# Pre-metastatic niche drives breast cancer invasion by modulating MSC homing and CAF differentiation

**DOI:** 10.1101/2021.01.12.426460

**Authors:** Neha Saxena, Garvit Bhardwaj, Sameer Jadhav, Hamim Zafar, Shamik Sen

## Abstract

The extent to which cancer-associated alterations in extracellular matrix stiffness influences the crosstalk between cancer cells and mesenchymal stem cells (MSCs) remains unclear. By analyzing multiple singlecell RNA sequencing datasets, we establish the existence of a cell sub-population co-expressing MSC and cancer associated fibroblast (CAF) markers in highly aggressive triple-negative breast cancers in primary tumor, secondary sites, and in circulatory tumor cell clusters. Using hydrogels of varying stiffness corresponding to different stages of cancer progression, we show that on pre-metastatic stroma mimetic 2 kPa gels, MDA-MB-231 breast cancer cell secreted conditioned media drives efficient MSC chemotaxis and induces stable CAF differentiation in a TGF*β*/contractility-dependent manner. In addition to enhancing cancer cell proliferation, 2 kPa CAFs maximally boost local invasion and confer resistance to flow-induced shear stresses. Together, our results suggest that homing of MSCs at the pre-metastatic stage and their differentiation into CAFs actively drives breast cancer invasion and metastasis.

## Introduction

Of the multiple types of cells present in the tumor microenvironment (TME), an accumulating body of evidence has demonstrated the importance of cancer associated fibroblasts (CAFs) in actively driving cancer progression by fostering tumor growth and invasion via matrix metalloproteinase (MMP) mediated remodeling of the extracellular matrix (ECM) [1–4]. While increased secretion of fibrillar collagen, fibronectin and laminin lead to stromal stiffening and desmoplasia, force-mediated remodeling of synthesized ECM leads to formation of linearized tracks thereby promoting cancer invasion [5]. Though most CAFs express the myofibroblast marker *α* smooth muscle actin (*αSMA*), considerable phenotypic heterogeneity exists within CAFs indicative of existence of multiple CAF sub-types with variable expression of fibroblast activation protein (*FAP*), fibroblast specific protein (*FSP*) and platelet derived growth factor receptor *β* (*PDGFRβ*) [6, 7]. In breast cancer, in comparison to less aggressive luminal sub-types, more aggressive HER2+ and triple negative breast cancers (TNBCs) have been shown to harbour CAFs exhibiting elevated levels of *FAP, αSMA, PDGFRβ* and *β*1 integrins [8].

Though the origin of CAFs remains debatable, several studies have identified mesenchymal stem cells as a potential source of CAFs [9–11]. Mesenchymal stem cells (MSCs) represent one of the prominent cell-types that home at the site of tumors in response to factors released by cancer cells [12, 13]. MSC homing has been documented in multiple types of cancers including breast, lung and pancreatic cancers [14–16]. Both MSCs as well as adipocyte derived stem cells (ASCs) are known to differentiate into CAFs upon exposure to cancer cell secreted transforming growth factor *β* (TGF*β*) [17]. MSCs, differentiated to CAFs, have been reported to increase the metastatic ability of weakly metastatic breast cancer cells with MSC secreted CCL5 implicated in inducing metastasis by enhancing cancer stem cell properties [9]. When co-cultured with breast cancer cells, MSCs have shown to exhibit higher homing through placental growth factor (PGF) and cancer metastasis through hypoxia inducible factor (HIF) [18]. In addition to secreting ECM proteins such as collagen and fibronectin [19], MSCs also play a crucial role in driving matrix stiffening through upregulation of lysyl oxidase (LOX) [18].

In epithelial cancers including breast, pancreatic and lung cancer, cancer aggressiveness and survival is associated with increased deposition, alignment and crosslinking of fibrillar collagens leading to several-fold increase in stromal stiffness [20, 21]. While MSCs are known to respond to matrix stiffness and differentiate into multiple lineages [22], the influence of stiffness in regulating MSC differentiation into CAFs is less understood. In a recent study, Keely and co-workers showed that upon exposure to cancer conditioned media (CM), MSCs differentiate into CAFs in a stiffness-dependent manner and drive cancer cell proliferation by secreting prosaposin [23]. However, the temporal dynamics of MSC homing at the site of tumors remains incompletely understood. Moreover, since stiffness regulates cancer cell secretome, it is likely that the response of MSCs to cancer CM is stiffness-dependent.

In this work, we address the importance of stiffness-dependent crosstalk between MSCs and MDA-MB-231 breast cancer cells in mediating MSC chemotaxis, MSC differentiation to CAFs and their subsequent involvement in cancer invasion. Based on analysis of single-cell RNAseq (scRNAseq) datasets from multiple breast cancer patients, we first show that aggressive triple negative breast cancers (TNBCs) harbour a subpopulation of cells exhibiting MSC/CAF markers. Using polyacrylamide gels of stiffnesses mimicking that of normal, pre-metastatic and metastatic stroma, we then show that MSC chemotaxis and CAF differentiation are optimal on pre-metastatic stroma-mimetic 2 kPa gels in a TGF*β*-dependent, contractility-dependent manner, with CAF differentiation stable on 2 kPa gels. Next, we illustrate the role of 2 kPa CAFs in promoting cancer cell proliferation, stromal invasion and in sustaining physiological shear stresses. Finally, we perform analysis of scRNAseq data from circulators tumor cells (CTCs) and metastatic tumors to establish the presence of MSC/CAF expressing cells in CTC clusters and secondary tumors. In addition to illustrating the prominent role of MSCs in mediating cancer metastasis via stiffness-dependent homing at the primary tumor and differentiation into CAFs, our results implicates the pre-metastatic niche as an active driver of cancer progression.

## Results

### Presence of a MSC/CAF signature in triple negative breast cancers (TNBCs)

Among the four molecular subtypes of breast cancer, the TNBC sub-type is the most aggressive subtype in terms of clinical course and is known to be characterized by extensive inter-tumor as well as intra-tumor heterogeneity [24]. The aggressiveness of TNBC is characterized by early onset of disease, higher incidence of metastasis, greater relapse rate and shorter overall survival [25]. In order to evaluate if the stromal component of TNBCs harbour MSCs that contribute to higher invasiveness observed in TNBC, we analyzed scRNAseq data from 551 cells that were sampled from 11 breast cancer patients [26]. These cells were divided into a group consisting of 214 cells from 5 TNBC patients and another group consisting of 337 cells that were sampled from patients with other molecular subtypes (2 luminal A, 1 luminal B and 3 HER2-positive patients). Within each group of cells, we identified transcriptionally distinct cell populations by performing clustering using Seurat [27] and subsequently tested the clusters for enrichment of known MSC markers including *CD90, CD106* [28] and *CD140b* [29]. While 3 clusters were identified in the group consisting of TNBC cells (Fig. 1A), a subset of 10 cells in cluster 1 (green) expressed the MSC markers (Fig. 1C). In contrast, cells from group 2 were divided into 7 cell types (Fig. 1B), none of which expressed the MSC markers (Fig. 1D).

**Figure 1:**
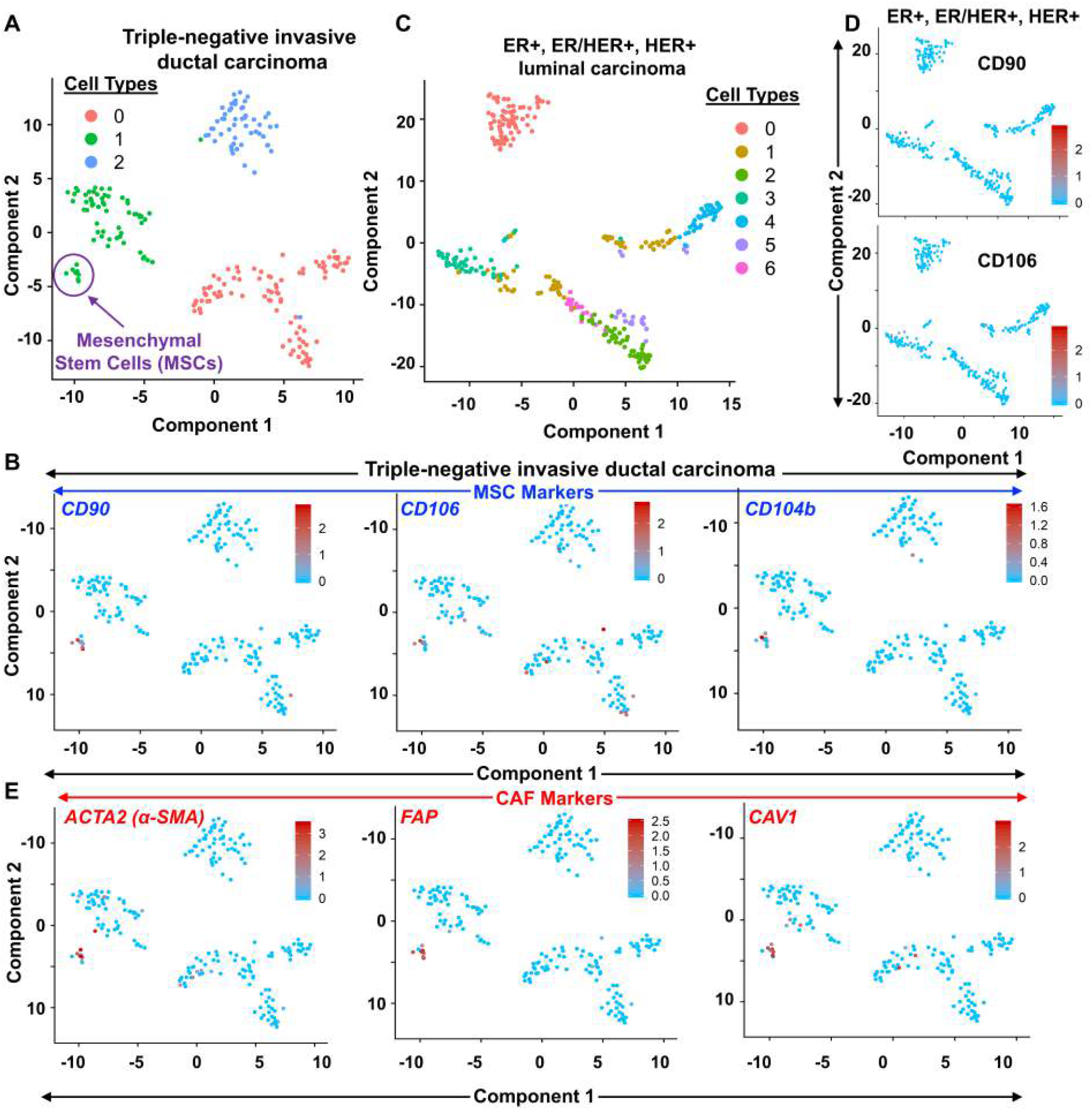
scRNAseq Analysis of TNBC and non-TNBC Data set: A) UMAP plot of cells from triple negative breast cancer (TNBC) patients delineates three cell type clusters, out of which cluster 1 harboured putative mesenchymal stem cells (MSCs). B) The subset of 10 cells in cluster 1 (green) expressed several MSC markers (CD90, CD106. CD104b). C) UMAP plot of cells from non-TNBC patients showing 7 cell type clusters. D) Cells from non-TNBC patients did not express MSC markers. E) The putative MSCs (Cluster 1) were also found to express known CAF markers, including α-SMA, FAP, CAV1.

For the TNBC group, we selected the 10 cells expressing MSC markers and performed differential gene expression analysis by comparing with all other TNBC cells (Supp. Fig. 1A). This analysis revealed *CD90 (THY1)* to be one of the top differentially expressed genes in these cells (Supp. Fig. 1B). In addition, other MSC markers including *COL1A1* [30], *COL1A2* [31], *POSTN* [32] were found to be differentially expressed in these 10 cells indicating these cells to be MSCs that invaded the tumor stroma. The putative MSCs identified from the scRNAseq dataset were further tested for the expression of CAF markers. The putative MSCs were found to express known CAF markers including *ACTA2* (also known as *αSMA*), *FAP, CAV1, CD29, TAGLN, COL1A2, DCN* in both the data sets (Fig. 1E, Supp. Fig. 1C) and these genes were further enriched in differential expression analysis (Supp. Figs. 1B). Along with the expression of CAF associated genes, the putative MSCs also express markers associated with invasion (*MMP2*), ECM stiffening (*LOX*) as well as markers associated with bone metastasis (*CTGF, IGF1*) (Supp. Fig. 1C).

We further checked another scRNAseq TNBC dataset for the presence of MSCs [33]. This dataset consisted of 24271 cells that were sampled from five primary TNBC patients. Clustering analysis with Seurat identified 20 cell type clusters of which three clusters (Clusters 6, 11 and 16) expressed different MSC markers (CD90, CD106, PDGFRB) (Supp. Fig. 2A). As before, MSC, CAF and invasion markers were found to be differentially expressed in the above three clusters (Supp. Figs. 2B, 3). To further characterize the lineage relationships between the cells expressing MSC/CAF signature in TNBC, 3 clusters were used from this dataset for trajectory inference. Differentiation trajectory of the cells in these 3 clusters was reconstructed using Monocle 3 [34], which inferred a branched trajectory consisting of three major branches (Supp. Fig. 4A). The cells were pseudotemporally ordered by selecting the cells expressing MSC markers *CD105* and *Stro-1* as the initial population in the trajectory. Cells in Branch 1 were the closest to the MSCs expressing the MSC marker CD90 as well as the CAF markers *ACTA2, CAV1* and *S100A4* (also referred to as *FSP1*) (Supp. Fig. 4B). These cells further differentiated into two branches, with Branch 2 expressing both MSC (*COL1A1, CD90*) and CAF markers (*ACTA2, DCN, COL1A2*) and Branch 3 expressing CAF markers (*FSP1, COL1A2, CAV1*). Collectively, the trajectory analysis suggests that the sub-population of cells expressing MSC markers in TNBC tissue differentiate into CAFs as well as cells that co-express CAF and MSC markers.

### Stiffness-modulated cancer conditioned media (CCM) regulates MSC chemotaxis

Based on the above anyalsis, we hypothesize that cells expressing MSC/CAF markers correspond to MSCs which home at the site of primary tumor and undergo partial differentiation into CAFs. Matrix stiffening is a hallmark of breast cancer progression [35–37]. To determine how biophysical alterations in ECM mechanics associated with cancer progression is temporally correlated with homing of MSCs at the site of primary tumor, we probed the influence of stiffness-dependent secretome of cancer cells in mediating MSC chemotaxis. Specifically, MSC chemotaxis was studied in response to factors secreted by MDA-MB-231 breast cancer cells cultured on polyacrylamide gels mimicking bulk stiffness of normal mammary stroma (0.5 kPa), pre-metastatic tumor stroma (≈ 2 kPa), and metastatic tumor stroma (≈ 5 kPa) [35, 36]. Chemotaxis experiments were performed using a microfluidic device comprising of two large side channels connected by several transverse channels wherein introduction of a chemokine in one of the side channels leads to establishment of a stable chemical gradient, leading to MSC chemotaxis (Fig. 2A) [38]. While MSCs mixed with collagen gels were seeded on one of the side channels in plain MSC culture media, the remaining portion was filled with only collagen gel (cell free side).

**Figure 2:**
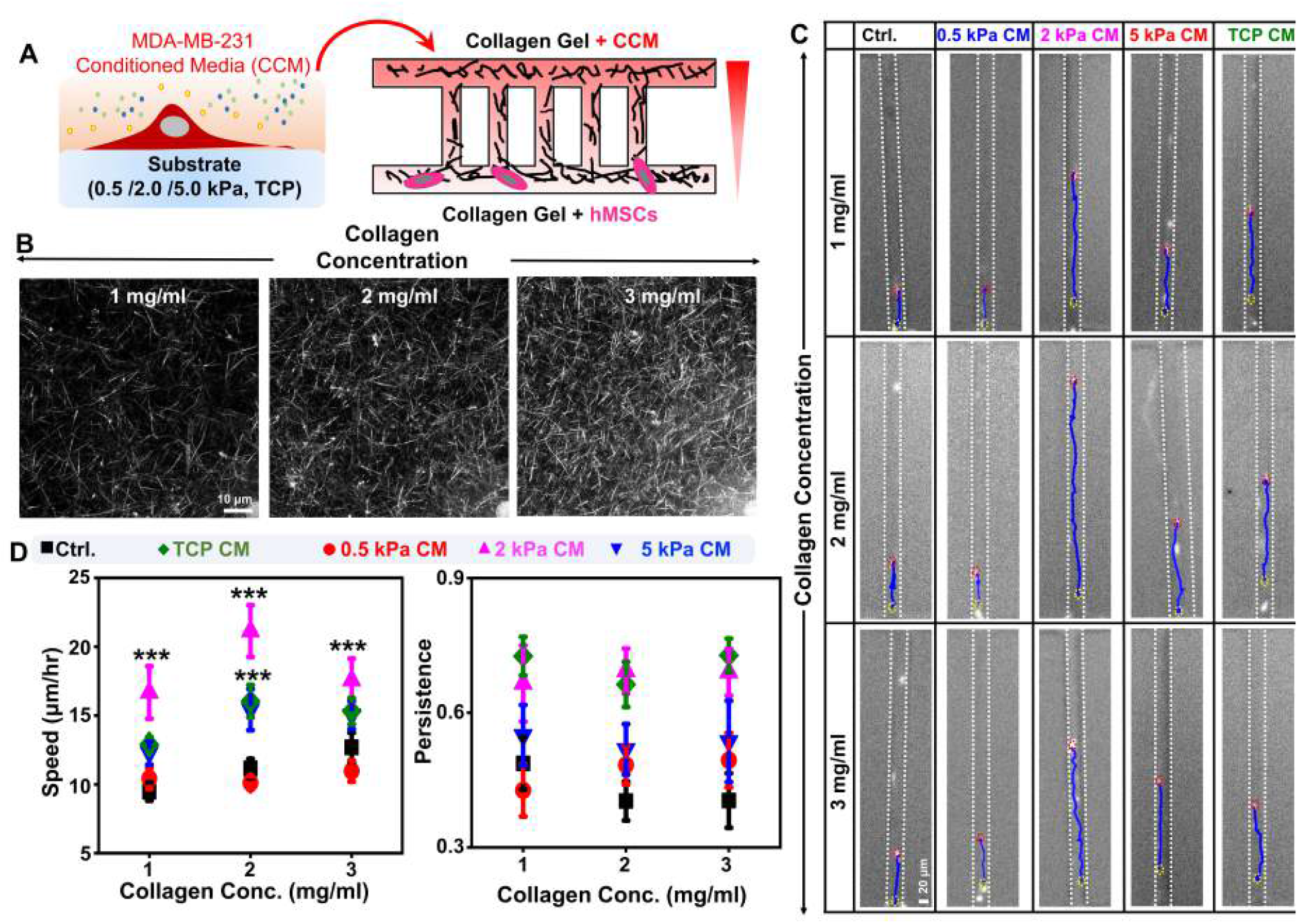
Influence of stiffness-modulated cancer cell conditioned media (CCM) on MSC chemotaxis. A) CCM collected from MDA-MB-231 breast cancer cells cultured on 0.5, 2, and 5 kPa polyacrylamide (PA) gels was used as a chemical cue to study MSC chemotaxis. Chemotaxis was studied within a microfluidic device consisting of two side-channels connected by multiple transverse channels. While MSCs mixed with 3D collagen solution was introduced in the bottom channel, collagen solution was introduced in the top channel. After collagen gel formation at 37°C, CCM was introduced in the top channel to setup a chemokine gradient within the transverse channels. B) Confocal reflectance images of polymerized collagen at different concentrations (1, 2, and 3 mg/ml). C) Representative trajectories (blue lines) of MSCs migrating through transverse channels in the presence or absence of stiffness-modulated CCM. Cells were tracked by labeling nuclei with Hoechst 33342. Scale bar = 20 *μ*m). D) Quantification of cell speed and persistence of MSCs migrating through transverse channels at varying collagen concentrations in the absence (Ctrl) and presence of CCM (*n* ≥ 30 cells per condition pooled from N = 3 independent experiments; error bars represent ± SEM; ∗ ∗ ∗ indicates statistical significance (*p* < 0.001) compared to 0.5 kPa CM condition).

Upon adhesion of MSCs to the surrounding collagen matrix, cancer conditioned media (CCM) collected from MDA-MB-231 cells cultured on gels (i.e., 0.5 kPa CM, 2 kPa CM and 5 kPa CM) and from tissue culture plates (i.e., TCPCM) was introduced at the cell free side. Experiments were performed by varying the concentration of collagen gels so as to generate matrices of varying pore sizes (Fig. 2B). Quantification of trajectories of individual MSCs (red lines) along the transverse channels (white dotted lines) tracked by labeling nuclei (blue) revealed marked differences in MSC motility across the different conditions (Fig. 2C). While cell motility was collectively influenced by collagen concentration and CCM composition, across all the collagen gels, fastest migration was observed in the presence of 2 kPa CM (Fig. 2D). Highest cell persistence was observed in the presence of 2 kPa CM and 5 kPa CM. Together, these results suggest that 2 kPa CM optimally induces MSC chemotaxis.

### Stiffness regulates MSC differentiation into CAFs in a TGF*β*-dependent manner

To next probe how stiffness of the tumor microenvironment influences the fate of MSCs which home at the site of primary tumors, MSCs were cultured for 7 days on 0.5 kPa, 2 kPa and 5 kPa gels using conditioned media (CM) composed of normal MSC media and CCM in 1:1 ratio (Fig. 3A). MSCs cultured on the respective gels in regular MSC media served as controls. While initial spreading at Day 1 was stiffness-dependent with CM making no difference, CM-incubated MSCs exhibited increased spreading on 2 and 5 kPa gels after 7 days in culture (Fig. 3B, C). In comparison, MSC motility was found to be sensitive to presence of CM on 2 kPa gels on Day 1 itself, with further increase in motility detected on 2 and 5 kPa gels at Day 7 (Fig. 3C). After 7 days in culture, in comparison to 0.5 kPa gels, cell spreading and motility were higher on 2 and 5 kPa gels in the presence of CM. Additionally, tracking of cell proliferation over a period of 7 days revealed highest proliferation on 2 kPa substrates (Supp. Figs. 5A, B).

**Figure 3:**
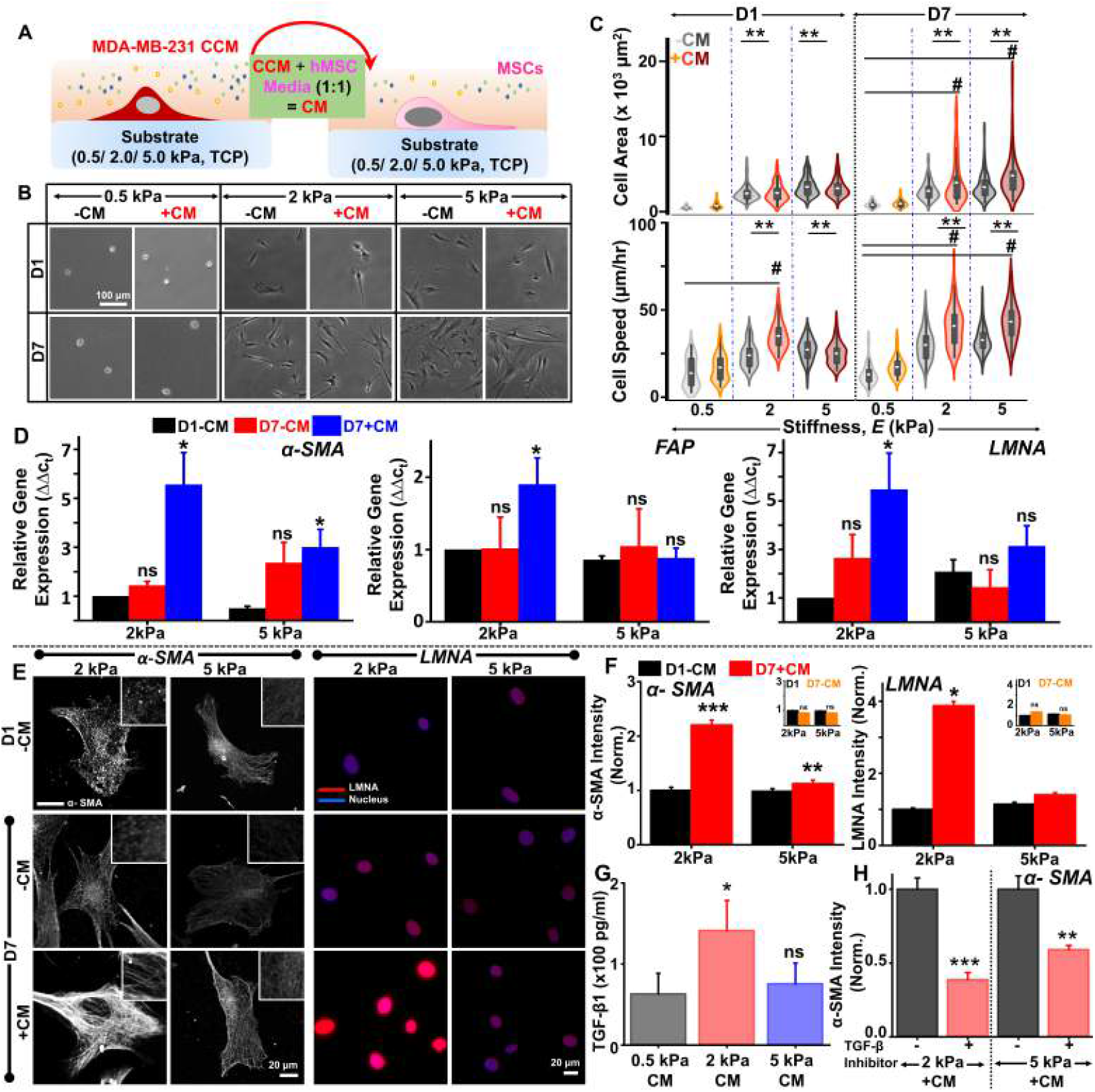
Matrix stiffness regulates MSC differentiation into CAFs: A) Experimental plan for studying the effect of stiffness-modulated CCM on MSC differentiation. MSCs were cultured in conditioned media (CM) generated by mixing stiffness-modulated CCM with MSC complete media in 1:1 ratio. B) Representative images of MSCs cultured on 0.5, 2, and 5 kPa PA gels ±CM. Images were acquired at Day 1 (D1) and Day 7 (D7). Scale bar = 100 μm. C) Quantification of MSC spreading and motility at D1 and D7 across different conditions (*n* ≥ 50 cells per condition pooled from N = 3 independent experiments; error bars represent ± SEM, ∗∗ signifies statistical significance (*p* < 0.01) w.r.t 0.5 kPa (-CM) condition at D1 and D7 respectively and # signifies statistical significance (*p* < 0.05) within subgroups). D) Expression profile of CAF markers (α-SMA, FAP) and differentiation marker (LMNA) in MSCs cultured on (2 and 5) kPa gels ±CM for 7 days (N = 3 independent experiments; Error bars represent ± SEM; ∗ signifies statistical significance (*p* < 0.05) w.r.t D1-CM condition). E, F) Representative images and intensity quantification of α-SMA/LMNA stained MSCs cultured on (2 and 5) kPa gels ±CM at D1 and D7. Scale bar = 20 μm. For image quantification, integrated intensities were normalized w.r.t 2 kPa, –CM, D1 condition (*n* ≥ 50 cells per condition from N = 3 independent experiments; error bars represent ± SEM, ∗ ∗ ∗, ∗∗, ∗ indicates statistical significance (*p* < 0.001, *p* < 0.01, *p* < 0.05, respectively) compared to D1-CM). Insets show integrated intensities normalized to 2 kPa, –CM, D1 condition for cells cultured in the absence of CM on (2 and 5) kPa gels. G) ELISA-based quantification of TGF*β* levels in stiffness-modulated CCM (N = 3 independent experiments; Error bars represent ± SEM; ∗ signifies statistical significance (*p* < 0.05) w.r.t D1-CM condition). Error bars represent ± SEM. H) α-*SMA* levels in MSCs cultured on (2 and 5) kPa gels ± TGF*β* neutralizing antibody for 7 days in the presence of CM (*n* ≥ 90 cells per condition pooled from N = 3 independent experiments; error bars represent ± SEM, ∗ ∗ ∗, ∗∗ indicates statistical significance (*p* < 0.001, *p* < 0.01, respectively) compared to no inhibitor condition).

Higher sensitivity of MSC spreading and motility on 2 kPa gels in the presence of CM correlated with increased cytoskeletal organization and higher cortical stiffness in these cells (Supp. Figs. 5C, D). Since expression of the MSC marker Stro-1 was downregulated after 1 week in the presence of CM on both 2 and 5 kPa gels (Supp. Figs. 6Ei, Eii), we hypothesized that the observed phenotypic changes were indicative of MSC differentiation into CAFs. Consistent with this, expression of CAF markers *α-SMA* and *FAP* [6, 7, 39] were elevated on 2 kPa gels in the presence of CM, but not on 5 kPa gels (Fig. 3D). Further, expression of the nuclear marker Lamin A (*LMNA*) associated with MSC differentiation increased 5-fold on 2 kPa, +CM condition (Fig. 3D). Consistent with these observations, immunostaining for *α-SMA* and *LMNA* revealed highest expression on 2 kPa, +CM condition (Figs. 3E, F).

Since TGF*β* is known to induce CAF differentiation [17], we first checked levels of TGF*β* in CCM collected from the different gels, with cell culture media used as blanks. Maximum upregulation of CAF markers on 2 kPa gels was associated with highest TGF*β* level on 2 kPa gels (Fig. 3G). To establish the importance of TGF*β* in driving CAF differentiation, experiments were performed wherein MSCs were incubated with CM in the presence of TGF*β* inhibitor (Supp. Fig. 6F). Antibody-mediated blocking of TGF*β* receptor led to ≥50% drop in *α*-SMA levels on both 2 and 5 kPa gels and ≈ 30% drop in *LMNA* levels on 2 kPa gels (Fig. 3H, Supp. Figs. 6Gi, Gii). Collectively, these results suggest that MSC differentiation to CAFs is driven by TGF*β* present in the CCM, with highest TGF*β* secreted by MDA-MB-231 cells on 2 kPa gels.

### CAF differentiation is contractility-dependent and is stable on 2 kPa gels

In line with highest *α*-SMA expression on 2 kPa gels, 2 kPa CAFs possessed highest pMLC levels in the presence of CM (Figs. 4A, B) and exerted highest tractions (Figs. 4C, D, Supp. Fig. 6H). Since stiffness-dependent differentiation requires non-muscle myosin II-dependent stiffness-sensing [22], to probe the role of contractility in mediating MSC differentiation into CAFs, experiments were performed wherein MSCs were cultured in ±CM in the presence of the non-muscle myosin II inhibitor blebblistatin (Bleb) for a period of 7 days (Fig. 4E). While Bleb treatment did not alter *α-SMA/LMNA* expression in the absence of CM, in cells cultured in the presence of CM, in line with significant reduction in pMLC levels in Bleb-treated cells (Supp. Figs. 6Ii, Iii), a dramatic drop in both *α-SMA* and *LMNA* intensities were observed (Figs. 4E, F). Together, these results highlight the role of contractility in mediating CM induced differentiation of MSCs into CAFs.

**Figure 4:**
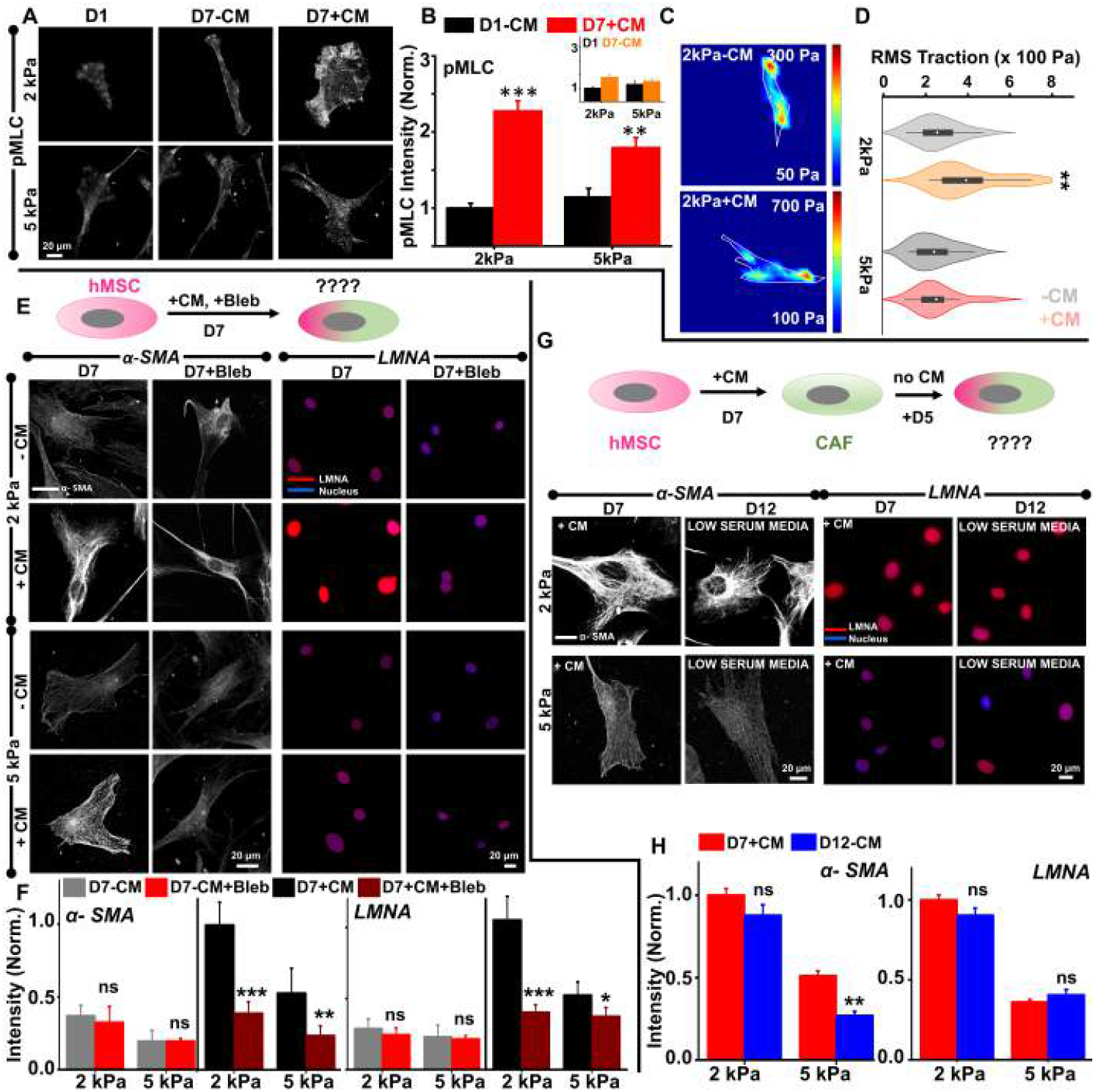
2 kPa CAFs are more contractile and exhibit stable differentiation: A, B) Representative images and intensity quantification of pMLC staining in MSCs cultured ±CM on (2 and 5) kPa gels for 7 days. Scale bar = 20 μm. For quantification of pMLC intensity, integrated intensities were normalized with respect to 2 kPa, −CM, D1 condition (*n* ≥ cells per condition pooled from N = 3 independent experiments; error bars represent ± SEM, ∗ ∗ ∗, ∗∗ indicates statistical significance (*p* < 0.001, *p* < 0.01, respectively) compared to D1-CM). Insets show integrated intensities normalized to 2 kPa, –CM, D1 condition for cells cultured in the absence of CM on (2 and 5) kPa gels. C, D) Representative traction force heatmaps and RMS traction force quantification in MSCs cultured ±CM on (2 and 5) kPa gels for 7 days (n = 12 cells per condition from N = 3 independent experiments; error bars represent ± SEM, ∗∗ indicates statistical significance compared to D1CM). E) To probe the effect of myosin inhibition on MSC differentiation into CAFs, MSCs were cultured in CM supplemented with 5 μM Blebbistatin (Bleb) for 7 days. Representative images and intensity quantification of α-SMA/LMNA stained MSCs cultured on (2 and 5) kPa gels for 7 days ±CM, ±Bleb. Scale bar = 20 μm. F) Integrated intensities were normalized with respect to 2 kPa, +CM condition (*n* ≥ 40 cells per condition pooled from N = 3 independent experiments; error bars represent ± SEM, ∗ ∗ ∗, ∗∗ indicates statistical significance (*p* < 0.001, *p* < 0.01, respectively) compared to D7+CM condition). G, H) Experimental plan to probe the stability of CAF differentiation. After 7 days culture in the presence of CM, MSCs were cultured on (2 and 5) kPa gels for another 5 days in complete media and stained for CAF markers. Representative images and intensity quantification of α-SMA/LMNA staining after day 5 of CM removal (i.e., Day 12 (D12)) on (2 and 5) kPa gels (Scale bar: 20 *μ*m). Integrated intensities were normalized with respect to 2 kPa, +CM, D7 condition (*n* ≥ 40 cells per condition pooled from N = 3 independent experiments; error bars represent ± SEM, ∗∗ indicates statistical significance (*p* < 0.01) compared to D7+CM condition).

To next assess the stability of MSC differentiation into CAFs, MSC-differentiated CAFs were cultured for another 5 days in culture in low serum media in the absence of CM (Fig. 4G). While the drop in *α-SMA/LMNA* levels on 2 kPa gels were marginal and statistically insignificant, *α-SMA* levels dropped significantly on 5 kPa gels (Fig. 4H). To further probe the sensitivity of CAF marker expression to contractile inhibition, CAFs were cultured for 3 days in the presence of Bleb in the absence of CM (Supp. Fig. 7A). Remarkably, levels of pMLC and α-SMA remained relatively unchanged on 2 kPa gels, but dropped significantly on 5 kPa gels (Supp. Figs. 7Bi, Bii). Collectively, these results suggest that 2 kPa substrates induce stable differentiation of MSCs into CAFs in a contractility-dependent manner.

### 2 kPa CAFs maximally enhance proliferation and invasiveness of cancer cells

To next probe the role of CAFs in modulating cancer cell behavior, MDA-MB-231 cells were cultured on 2 and 5 kPa gels in the presence of CAF secreted media, i.e., CAF conditioned media (CAFCM) (Fig. 5A). While CAFCM was found to induce increased cell spreading on both the gels (Figs. 5B, C), tracking of cell proliferation over a 7 day time period revealed fastest proliferation on 2 kPa gels supplemented with CAFCM (Figs. 5D, E).

**Figure 5:**
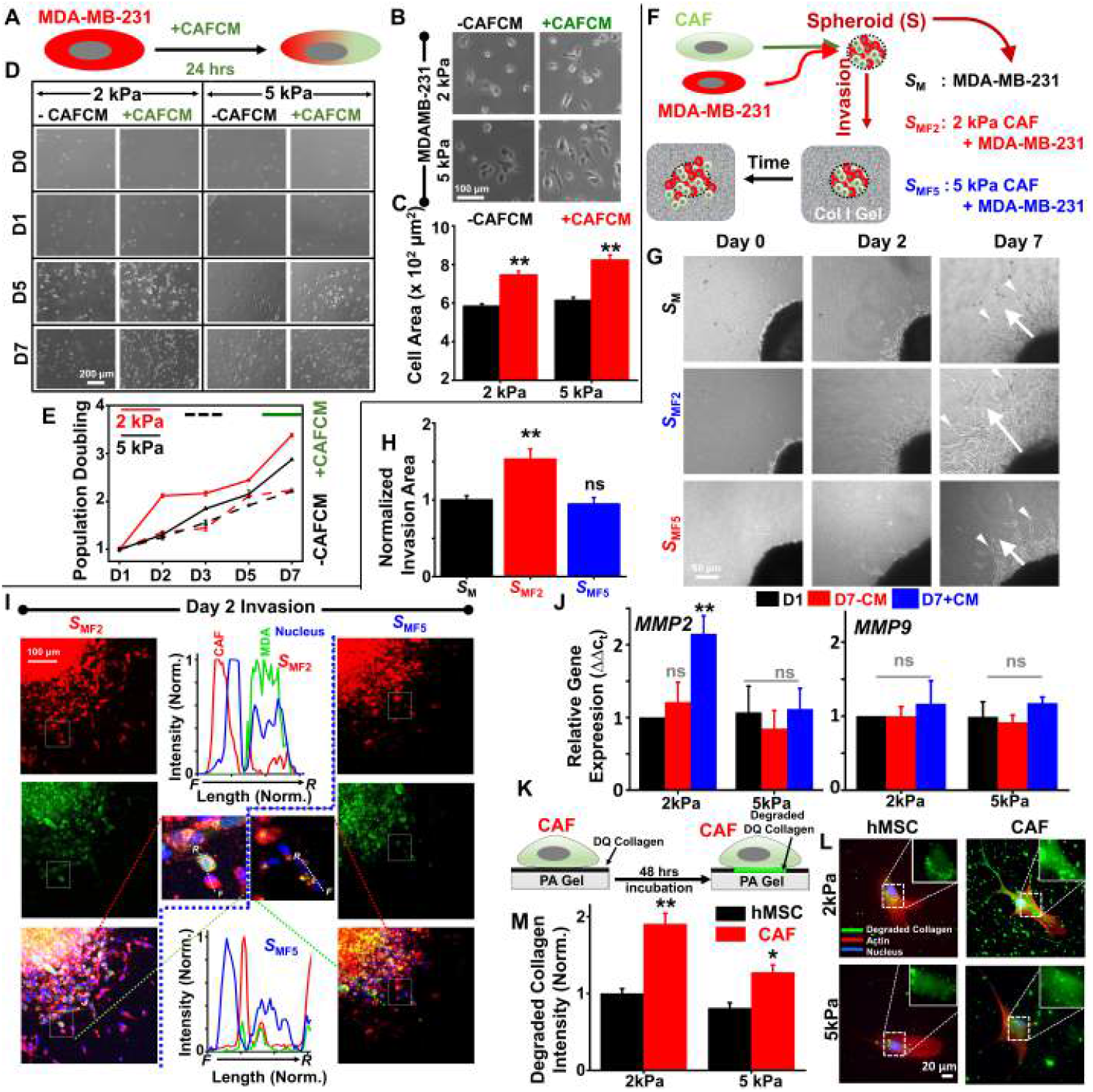
2 kPa CAFs maximally enhance proliferation and invasiveness of MDA-MB-231 breast cancer cells: A) Cancer cells were cultured in stiffness-modulated CAF secreted CM (CAFCM) for 24 hrs. B) Representative images of MDA-MB-231 cultured on (2 and 5) kPa gels ±CAFCM. Scale bar = 100 μm. C) Quantification of MDA-MB-231 cell spread area ±CAFCM (*n* ≥ 50 cells per condition pooled from N = 3 independent experiments; error bars represent ± SEM, ∗∗ indicates statistical significance (*p* < 0.01) compared to –CM). D) Representative images of MDA-MB-231 cultured on (2 and 5) kPa gels ±CAFCM for 7 days. Scale bar = 200 μm. E) Quantification of population doubling of cancer cells ±CAFCM (N = 3 independent experiments; error bars represent ± SEM). F) Schematic of spheroid formation using cancer cells alone (SM) and by combining cancer cells with (2 and 5) kPa CAFs (SMF 2 and SMF 5, respectively). Spheroid invasion was performed by embedding spheroids in 3D collagen gels. G) Representative temporal images of invasion by SM, SMF2 and SMF5 spheroids in 3D collagen gels. Scale bar = 50 μm. White arrows depict the extent of outward invasion. H) Quantification of area invaded by SMF 2 and SMF 5 spheroids normalized to that of SM spheroids (N = 3 independent experiments; error bars represent ± SEM; ∗∗ indicates statistical significance (*p* < 0.01) compared to SM, ns indicates not significant). I) Spatial positioning of CAFs and MDA-MB-231 cells at the invasive front. CAFs were stained with CellTracker red and MDA-MB-231 with CellTracker green. Intensity profiles along lines drawn in the boxed regions show the relative positions of CAFs and MDA-MB-231 cells from front (F) to rear (R). J) Expression profile of matrix degrading enzymes MMP2 and MMP9 in MSCs cultured on (2 and 5) kPa gels ±CM for 7 days (N = 3 independent experiments, ∗∗ indicates statistical significance (*p* < 0.01) compared to D1, ns indicates not significant). K) Schematic of assessing MMP mediated matrix degradation. Cells were plated on gels functionalized with collagen mixed with DQ collagen. MMP mediated degradation was visualized by fluorescence signal. L, M) Representative images and intensity quantification of degraded collagen by MSCs on 2 kPa/5 kPa gels and by 2 kPa/5kPa CAFs. Integrated intensities were normalized w.r.t 2 kPa D1 MSCs (*n* ≥ 55 cells per condition pooled from N = 3 independent experiments; error bars represent ± SEM, ∗∗, ∗ indicates statistical significance (P < 0.01, 0.05) w.r.t D1 MSCs.)

To next investigate the role of CAFs in regulating cancer invasion, spheroids (*S*) were prepared using hanging drop method with MDA-MB-231 cells alone (S_M_) or by combining MDA-MB-231 cells with 2 kPa CAFs or 5 kPa CAFs (S_MF2_ and S_MF5_, respectively) (Fig. 5F). The sizes of the three different types of spheroids were similar with most cells within the spheroids being viable (Supp. Figs. 7C-E). Invasiveness of spheroids was assessed by implanting single spheroids in 3D collagen gels and tracking the extent of outward cell scattering over a period of 7 days (Fig. 5G). Quantification of the area invaded by cells migrating out from the spheroids revealed maximum invasiveness of spheroids formed from 2 kPa CAFs, i.e., S_MF2_ spheroids (Fig. 5H).

Recent literature has demonstrated the role of CAFs in driving collective invasion of cancer cells [40]. To probe the role of 2 kPa and 5 kPa CAFs in leading cancer invasion, CAFs were stained with CellTracker Red and MDA-MB-231 with CellTracker Green to visualize the spatial position of the two cell types within the spheroids and also during the outward invasion. Within the heterospheroids implanted in collagen gels (i.e., Day 0 spheroids), the extent of CAF enrichment at the outer periphery was clearly observed in SMF2 spheroids, but not in SMF5 spheroids (Supp. Fig. 7F). Consistent with this, while the invasive front of SMF2 spheroids comprised primarily of 2 kPa CAFs, the invasive front of SMF2 spheroids were populated by both 5 kPA CAFs as well as MDA-MB-231 cells (Fig. 5I). Highest invasiveness of SMF2 spheroids was associated with highest expression of MMP2 in 2 kPa CAFs, with MMP9 expression remaining unaltered (Fig. 5J). In line with highest MMP2 expression, quantification of the extent of collagen degradation visualized using DQ collagen revealed maximum degradation (i.e., fluorescence) by 2 kPa CAFs (Figs. 5K-M). Taken together, these results suggest that in addition to promoting cancer cell proliferation, 2 kPa CAFs drive cancer invasion via increased matrix degradation.

### Spheroids formed using 2 kPa CAFs are more adhesive and resistant to shear stresses

Thus far, our results suggest that 2 kPa CAFs enhance stromal invasiveness of cancer cells. Cancer cell metastasis involves entry into the vasculature, adhesion to the endothelium and subsequent extravasation. To probe the possible role of CAFs in modulating adhesion and shear resistance of heterospheroids, integrin profiling of CAFs and adhesivity of heterospheroids were performed. Integrin profiling of CAFs revealed significant upregulation in expression of β1 and β3 integrins in 2 kPa CAFs, but not in 5 kPa CAFs (Figs. 6A, Bi, Bii). To next probe the consequence of increased integrin expression in 2 kPa CAFs in modulating adhesion of cancer cells, adhesion rate and adhesion strength of SM, SMF2 and SMF5 spheroids were assessed. For adhesion experiments, spheroids allowed to attach on to fibronectin-coated coverslips were washed after 1 hr to check the proportion of spheroids that remained attached (Fig. 6C). While the proportion of attached spheroids was comparable between SM and SMF5 spheroids, the number was higher for SMF2 spheroids (Figs. 6Di, Dii). To assess the strength of adhesion, spheroids were allowed to attach onto fibronectin-coated coverslips for 2 hrs, and then incubated with warm trypsin to assess the time required for detachment (τ_detach_) (Fig. 6E), with the exact time of detachment determined based on movement of spheroid centroids from their initial positions (Fig. 6Fi). The detachment duration, τ_detach_, given by the expression τ_detach_ = (τ ^*^ – 120) min, serves as a surrogate for adhesion strength with higher values indicative of stronger adhesion. Quantification of detachment timescales revealed that SMF2 spheroids exhibited stronger adhesion and took nearly twice as long to detach in comparison to SM and SMF5 spheroids (Fig. 6Fii).

**Figure 6:**
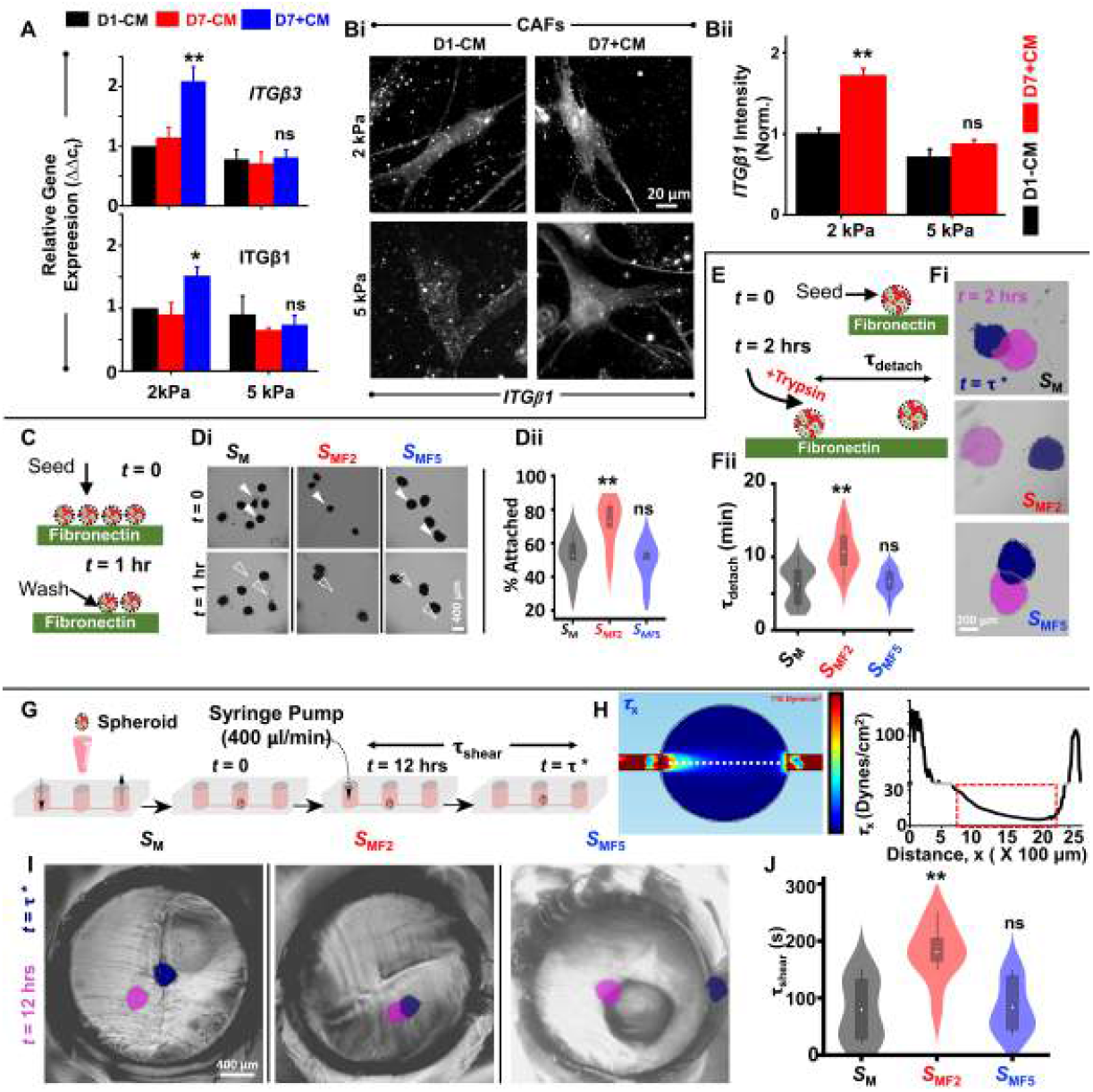
2 kPa heterospheroids are maximally adhesive and resistant to shear stresses: A) Expression profile of integrin β1 (ITGB1) and integrin β3 (ITGB3) in MSCs cultured on (2 and 5) kPa gels ±CM for 7 days (N = 3 independent experiments; error bars represent ±SEM, ∗∗, ∗ indicate statistical significance (*p* < 0.01, *p* < 0.05) compared to D1-CM, ns indicates not significant). Bi, ii) Immunostaining and quantification of ITGB1 in MSCs (D1-CM) and CAFs (D7+CM) on 2 and 5 kPa gels. Integrated intensities were normalized w.r.t 2 kPa D1 MSCs (*n* ≥ 55 cells per condition pooled from N = 3 independent experiments; error bars represent ± SEM, ∗∗ indicates statistical significance (*p* < 0.01) w.r.t D1-CM, ns indicates not significant.) C) Schematic of adhesion assay. SM, SMF2 and SMF5 spheroids were seeded on fibronectin (FN) coated coverslips and allowed to adhere for t = 1 hr. After removing loosely attached spheroids by washing with PBS, samples were imaged. Di) Representative images of spheroids at t = 0 hr and t = 1 hr. Scale bar = 400 μm. Dii) Percentage of SM, SMF2 and SMF5 spheroids that remained attached (*n* ≥ 53 spheroids per condition pooled from N = 3 independent experiments; error bars represent ± SEM; ∗∗ indicates statistical significance (*p* < 0.01) w.r.t SM spheroids, ns indicates not significant.). E) Schematic of de-adhesion assay. Spheroids seeded on FN coated coverslips were allowed to adhere for 2 hrs. After 2 hrs, trypsin was added to coverslips and spheroids were imaged till they were dislodged at time t = τ ^∗^(i.e., moved from their initial position). τdetach = t – 120 represents the duration (in minutes) required for a spheroid to de-adhere, and is a measure of adhesion strength. Fi, ii) Representative images of spheroids at t = 2 hrs (pink) and at t = τ ^∗^(blue) and quantification of τdetach (*n* ≥ 14 spheroids per condition pooled from N = 3 independent experiments; error bars represent ± SEM; ∗∗ indicates statistical significance (*p* < 0.01) w.r.t SM spheroids, ns indicates not significant). Scale bar = 200 μm. G) Experimental setup for assessing adhesion strength of individual spheroids under shear stresses. Spheroids were seeded in the middle port of a straight channel and allowed to attach for 10-12 hrs. Subsequently, spheroids were sub jected to a flow rate of 400 μl/min using a mechanical syringe pump until the spheroids detached from the substrate. H) Spatial map of shear stresses generated in the middle port of the device simulated in COMSOL using laminar flow physics. Graph represents variation of shear stress along the white dotted line. I) Representative images of spheroids after adhesion at t = 12 hrs (pink) and at the time of detachment (t = τ ^∗^, blue). Scale bar = 200 μm. J) Quantification of time required for detachment of spheroids under shear (τshear) (*n* ≥ 7 spheroids per condition pooled from N = 3 independent experiments; error bars represent ± SEM; ∗∗ indicates statistical significance (*p* < 0.01) w.r.t SM, ns indicates not significant).

To further test if SMF2 spheroids–which exhibit faster attachment and stronger adhesion–also exhibit greater resistance to shear, shear flow experiments were performed using a straight microfluidic channel with an inlet port, an outlet port, and an intermediate port for introducing spheroids (which was sealed after introducing spheroids through it) (Fig. 6G). After allowing the spheroids to attach onto the fibronectin-coated base of the microfluidic channel for 8 hrs, the inlet port was connected to a syringe pump. For a physiologically relevant flow rate of 400 μl/min [41], simulations predicted buildup of shear stresses upto 20 Pa (Fig. 6H). Plotting of shear stresses along the dotted white lines suggest spheroids situated at the middle of the long channel (x = 400 *μ*m) are subjected to shear stresses of ≈ 20 Dynes/cm^2^. Consistent with longer trypsin de-adhesion timescales, the time taken by spheroids to detach under shear, i.e., τ_shear_ –indicative of strength of adhesion–was nearly twice as higher for SMF2 spheroids compared to that of SM and SMF5 spheroids (Figs. 6I, J). Collectively, these results suggest that increased integrin expression on 2 kPa CAFs makes SMF2 spheroids more adhesive and resistant to shear stresses.

### Presence of MSCs/CAFs in circulating tumor cell (CTC) clusters and in secondary metastases

Cancer metastasis is mediated by clusters of circulating tumor cells (CTCs) with CTC clusters possess 50-fold higher chance of initiating metastasis compared to individual CTCs. To evaluate if MSCs and/or CAFs are present in CTC clusters, we analyzed scRNAseq data from CTCs isolated from invasive breast cancer patients and MMTV-PyMT mouse models [42]. The dataset consisted of 61 CTCs which were preprocessed and clustered into distinct cell types using Seurat. We identified three CTC clusters each of which expressed MSC markers CD29, CD73, CD166 (all clusters), CD105 (clusters 0 and 1), VCAM1 (cluster 0) and CD45 (cluster 1). The clusters also showed enrichment of CAF markers including ACTA2, FSP1, CAV1 and TAGLN. Apart from these, the CTCs also expressed CD44, LMNA, MMP9 as well as the bone metastasis marker CTGF (Fig. 7A and Supp. Fig. 8A). This suggests the presence of cell types other than cancer cells in CTC clusters which might help them to survive in blood flow and effectively metastasize to secondary sites.

**Figure 7:**
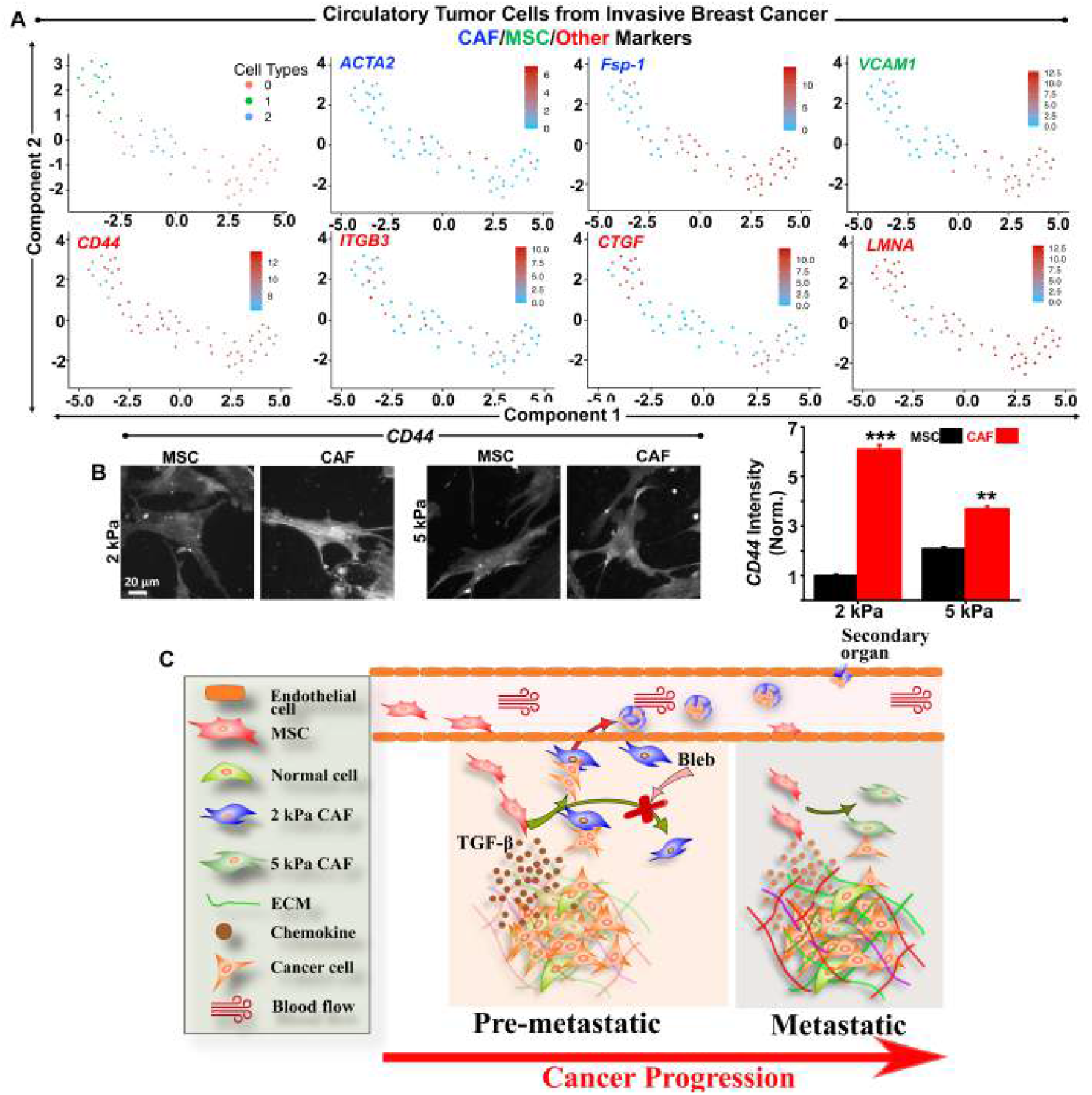
Presence of CAFs in circulating tumor cell (CTC) clusters and proposed model of pre-metastatic niche as a driver of cancer progression: A) scRNA sequencing from invasive breast cancer patients and MMTV-PyMT mouse mmodels. 61 CTCs were clustered into 3 cell types. Each cell cluster expressed MSC and CAF markers. Other than these markers CTCs also expressed CD44 (Cohesion), ITGB3 (Adhesion), LMNA (Differentiation) and CTGF (Bone metastasis). B) Immunostaining and quantification of CD44 in MSCs (D1-CM) and CAFs (D7+CM) on 2 and 5 kPa gels. Integrated intensities were normalized with respect to 2 kPa D1 MSCs (*n* ≥ 55 cells per condition pooled from N = 3 independent experiments; error bars represent ± SEM, ∗ ∗ ∗, ∗∗ indicate statistical significance (*p* < 0.001, *p* < 0.01, respectively) w.r.t D1-CM). C) Factors secreted by cancer cells on pre-metastatic stroma mimetic 2 kPa gels optimally induce MSC recruitment at the site of tumors and differentiation into CAFs. In addition to driving cancer cell proliferation through secreted soluble factors, differentiated CAFs aid in stromal invasion as well as distant metastasis by increasing adhesivity of cancer cells.

CD44-mediated homophilic interactions have been shown to drive multicellular aggregation in TNBCs [43]. Given the prominent expression of CD44 in all the three CTC clusters, and the presence of cells exhibiting MSC/CAF marker expression, we next checked CD44 expression in 2 kPa and 5 kPa CAFs. In comparison to baseline CD44 expression in MSCs, CD44 expression increased significantly in both 2 and 5 kPa CAFs with highest expression in 2 kPa CAFs (Fig. 7B). Though CD44 expression in cancer cells was found to be 10-15 fold higher compared to CAFs (Supp. Figs. 7Gi, Gii), staining of SMF2 spheroids invading collagen gels revealed comparable levels of CD44 at the interface of CAFs and MDA-MB-231 cells in contact at the invasive front (Supp. Fig. 7H).

To finally test if CTCs expressing MSC/CAF signature were associated with metastasis, we integrated single-cell RNA sequencing data from primary and metastatic tumor cells (375 cells in total) captured during seeding of micrometastasis in patient-derived-xenograft models of breast cancer [44] with the 61 CTC scRNAseq data [42]. To eliminate batch effects, we performed anchor-based integration using Seurat [45] and clustered the cells. The cells were clustered into two groups out of which the larger cluster (cluster 0) consisted of 54 CTCs, 121 primary and 129 metastatic tumor cells (Supp. Fig. 8B), whereas the other cluster only contained 7 CTCs and was dominated by 71 primary and 54 metastatic cells. The cluster harbouring both CTCs and metastatic cells in larger proportion (i.e., cluster 0) showed enrichment for MSC markers (CD73), CAF markers (ACTA2, CAV1) and other markers (CD44 and LMNA). This analysis suggests that CTCs that express MSC/CAF signature were also associated with primary and metastatic cells expressing the same markers.

## Discussion

In this study, by analyzing publicly available scRNAseq data from breast cancer patients, we first correlated the aggressiveness of TNBCs with presence of a sub-population of cells exhibiting MSC/CAF signature. We then showed that stiffness-modulated CCM regulates MSC chemotaxis and differentiation into CAFs. Next, we demonstrated the enhanced capacity of 2 kPa CAFs in enhancing invasiveness of cancer cell and imparting resistance to flow-induced shear stresses. Finally, by analyzing scRNAseq data from CTCs, we showed that CTC clusters harbour cells exhibiting MSC/CAF markers. Based on our findings, we propose a model wherein cancer progression is associated with MSC homing to tumors at the pre-metastatic stage, differentiation into CAFs, and subsequent CAF driven invasion and metastasis (Fig. 7C). Our results suggest that the combination of physical and chemical cues present in the pre-metastatic niche actively drives cancer progression by efficiently recruiting MSCs and stably differentiating them into CAFs. In addition to driving local invasion, our results suggest that MSC differentiated CAFs may aid in secondary organ colonization through increased adhesion under shear.

The tumor microenvironment (TME) represents a dynamic environment undergoing continuous alterations that collectively drive cancer aggressiveness. In addition to dynamic alterations in ECM properties, the presence of of multiple cell types including immune cells, resident fibroblasts, pericytes, CAFs as well as MSCs highlight the cell level heterogeneity within the TME [8, 46]. By analyzing publicly available scRNAseq datasets, we have established the presence of cells co-expressing MSC/CAF makers in TNBCs, but not in other subtypes of breast cancer. Trajectory analysis suggests that MSC differentiation can lead to generation of CAFs as well as an intermediate differentiated phenotype where both MSC and CAF markers are co-expressed. However, the physical cues determining these differentiation trajectories remain unclear. By probing stiffness-mediated bi-directional crosstalk between MSCs and cancer cells, we establish the importance of the pre-metastatic niche in giving rise to this mixed MSC/CAF signature. In our experiments, conditioned media secreted by cancer cells on pre-metastatic stroma-mimetic 2 kPa gels optimally induced MSC chemotaxis. MSCs are known to migrate in response to several tumor site-specific soluble factors including interleukin-6 (IL-6), IL-1, TGF*β* 1, epidermal growth factor (EGF) and stromal derived factor 1 (SDF-1) [12, 47]. Fastest MSC chemotaxis induced by 2 kPa cancer conditioned media (CCM) may be partly attributed to elevated TGF*β* levels. In a recent study on stiffness-dependent modulation of secretome of breast cancer cells, sphingosine-1-phosphate (S1P) was identified as an important bioactive lipid secreted by MDA-MB-231 cells to a greater extent on softer substrates [48]. Interestingly, S1P is a potent chemoattractant for MSCs driving their exit from the bone marrow to the peripheral blood [49]. Thus, increased MSC chemotaxis in response to 2 kPa CCM may be driven by the presence of multiple factors in the CCM and warrant a detailed proteomic analysis.

CAFs represent one of the most abundant cell types in the TME and constitute a heterogeneous population of mesenchymal-origin cells exhibiting both tumor-promoting and tumor-suppressive effects. Microenvironmental heterogeneity between low grade and high grade tumors has been shown to induce formation of different subtype of CAFs [50]. In breast cancer, four different CAF subsets have been identified based on expression of integrin β 1, α-SMA, FSP1, FAP, PDGFR and CAV1 [8]. In a recent study, Keely and co-workers demonstrated that CCM induces stiffness-dependent MSC differentiation into CAFs expressing high α-SMA [23]. However, the CCM used in this study was collected from cancer cells cultured on tissue culture plastic (TCP). Here, we show that stiffness-dependent CCM directly influences the expression profile of differentiated CAFs with a α-SMA^hi^FAP^hi^signature observed on 2 kPa gels. Our findings are consistent with a recent study wherein ECM composition and stiffness were identified as factors regulating TGF*β*-induced differentiation of murine pulmonary fibroblasts into CAFs with FAP^hi^ expressing CAFs observed on soft substrates [51]. FAP is a postprolyl peptidase capable of degrading both collagen and gelatin, and is involved in matrix turnover [52]. High FAP expression has been correlated with distant metastasis and poor prognosis in several solid tumors including breast cancer [37, 53–55]. MSC differentiation into FAP^hi^ CAFs on 2 kPa gels thus highlights the role of the pre-metastatic niche in actively driving cancer progression.

Cancer cell secreted TGF*β* leads to the conversion of stromal fibroblasts into myofibroblastic CAF subtype by inducing expression of α-SMA and FAP [3]. Consequently, blocking TGF*β* in-vivo inhibits the formation of this myofibroblastic CAF subtype [56]. Highest α-SMA and LMNA expression on 2 kPa gels coincided with highest levels of TGF*β* secreted on these gels. In line with this drop in α-SMA levels upon TGF*β* receptor blocking highlights the role of stiffness-dependent TGF*β* signaling in mediating CAF differentiation. TGF*β* is known to activate Rho/ROCK signaling pathway and induces expression of actin associated proteins such as calgizzarin, cofilin, and profillin [57]. Increased expression of α-SMA participates in the formation of stress fibres, and contributes to increased cytoskeletal tension and contractile properties of CAFs [57, 58]. In line with this, 2 kPa CAFs were found to possess highest levels of activated myosin (pMLC) and were the most contractile. Bleb-mediated inhibition of actomyosin contractility illustrates the importance of stiffness-sensing for MSC differentiation into CAFs. In addition, Bleb treatment has also been shown to inhibit conversion of latent TGF*β* to its active form which is required for CAF differentiation [59, 60]. CAFs have been shown to exhibit plasticity, i.e., they exhibit inter-conversion from one state to another [61, 62], may depend on the extent of differentiation. Highest LMNA expression on 2 kPa gels combined with relative insensitivity of 2 kPa CAFs to Bleb treatment is indicative of stable CAF differentiation on 2 kPa gels.

In comparison to 5 kPa CAFs, 2 kPa CAFs led to increased cancer cell proliferation and invasion. Increased proliferation of breast cancer cells may be driven by several CAF secreted factors including SDF-1, FGF1 and uPA [63]. Recently, Keely and co-workers showed that prosaponin secreted by MSCs on stiff substrates promotes proliferation of 4T1 mouse mammary cancer cells via AKT activation [23]. In contrast to this study, higher cancer cell proliferation observed on 2 kPa gels compared to 5 kPa gels may be attributed to the use of CAFCM instead of MSCCM. Maximal invasiveness of 2 kPa CAF-containing heterospheroids reported in our study may be attributed to the combination of higher *MMP/α-SMA* expression and increased contractility of 2 kPa CAFs leading to a combination proteolytic and force-driven remodeling of the ECM. In vivo, αSMA^hi^FAP^hi^CAFs drive cancer progression by LOX-mediated matrix stiffening and alignment generating matrices amenable for cancer invasion [64, 65]. In addition to path generation, CAFs have also been reported to physically pull out cancer cells from tumors through formation of heterotypic adhesions [40]. *CD44* + fibroblasts have been shown to promote survival and drug resistance of cancer cells [66]. Given the ability of *CD44* to form homophilic adhesions [43], it is possible that *CD44* also mediates adhesion between CAFs and MDA-MB-231 cells.

Secondary metastasis is mediated by CTC clusters which circulate in the blood stream, adhere to the endothelium and subsequently extravasate to seed secondary tumors, with CTC clusters possessing increased metastatic potential compared to single CTCs [67]. In addition to mediating stromal invasion, recent studies have documented the presence of CAFs in CTC clusters [68, 69]. Our analysis of scRNAseq data also suggests the presence of CAFs in CTC clusters in breast cancer patients. To overcome the shear stress from blood flow, cancer cells are known to associate with platelets [70]. In a recent study, King and co-workers demonstrated that CAFs confer fluid shear stress resistance to prostate cancer cells via intercellular contacts thereby maintaining viability and proliferative capacity of cancer cells [71]. Our results of increased adhesion and shear resistance of SMF 2 spheroids are consistent with this study. The prominent expression of CD44 observed in CTC clusters may not only contribute to maintaining cohesion of CTC clusters, but also mediate endothelial adhesion [72]. In addition to CD44, endothelial adhesion may also be mediated by β 1/β 3 integrins via binding to the fibronectin on the surface of endothelial cells [73, 74].

Different tumor types exhibit different secondary organ preference during metastasis. Consequently, identifying factors driving organ-specific metastasis represents an area of active research. Primary breast tumors are capable of metastasizing to several organs including lungs, liver, brain and bone, with bone frequently being the preferred site [75]. The presence of fenestrated endothelial cells and a discontinuous basal lamina in the bone marrow make it relatively easier for CTC clusters to seed bone metastasis [76]. In a seminal study, Massagúe and co-workers also demonstrated that MSC enriched triple negative breast tumors exhibit bone metastasis with MSC differentiated CAFs secreting high levels of CXCL12 and IGF1 [77]. Consistent with this, CTGF [78] and IGF1 [79], associated with bone metastasis, were also observed in the cells expressing MSC/CAF markers within TNBC tumors. These factors which are present at high levels in the bone marrow, in turn lead to establishment of clones with high Src activation and Src dependent PI3-Akt activation which increases the preference of breast cancer cells for the bone environment [77]. In addition, bone colonization may be mediated by β 3 integrins expressed both by CAFs and MDA-MB-231 cells, that can bind to osteopontin and vitronectin expressed by osteoblasts [80].

In conclusion, our study illustrates how stiffness-dependent crosstalk between MSCs and breast cancer cells drives cancer invasion through increased MSC homing, CAF differentiation and subsequent CAF-mediated invasion. Future studies targeting the crosstalk between MSCs and cancer cells at the pre-metastatic stage may lead to identification of therapeutic strategies for delaying disease progression.

## Materials & Methods

### scRNAseq data processing, clustering and visualization

We first preprocessed the scRNAseq datasets by filtering out cells that have unique feature counts over 2500 or less than 200. The cells containing > 5% mitochondrial counts were further removed. The cells remaining after the pre-processing steps were subject to normalization, and scaling using the Seurat V3 package [27]. For clustering the cells, we first selected the top variable genes (2000) using the FindVariableFeatures function in Seurat. Principal component analysis (PCA) was performed for reducing the dimensions of gene-expression data to the top 20 principal components. For visualizing the cells, the PCA space representation of the cells were projected to 2D using Uniform Manifold Approximation and Projection (UMAP) algorithm. The FindNeighbors and FindClusters in Seurat V3 were used for clustering the cells on PCA space using the Shared Nearest Neighbor (SNN) algorithm. The resulting cell clusters were visualized on UMAP space using the DimPlot function in Seurat. The expression of marker genes on the clusters were visualized using FeaturePlot function. The differentially expressed genes in each cluster were inferred using the FindAllMarkers function in Seurat. DoHeatMap function was used for generating the expression heatmap for cell clusters and top 20 differentially expressed genes for all clusters.

### Trajectory inference and pseudotemporal ordering

The differentiation trajectory of the cells expressing MSC/CAF markers was reconstructed using Monocle 3 [34]. In order to reconstruct the trajectory, first the selected cells were projected to a lower dimension using the reduceDimension() function in Monocle 3 which uses UMAP underneath. Next, the cells were partitioned using cluster cells() function to look for potentially discontinuous trajectories. In our case, all the cells were grouped into 1 cluster. The learn graph() function was used for learning a principal graph that delineates the possible trajectory of the cells. The pseudotime for the cells were calculated using the order cells() function by selecting the cells expressing MSC markers (CD105 and Stro-1) as the initial population. The expression of the marker genes on the trajectory was visualized using the plot cells() function.

### Integration of scRNAseq datasets

For integrating the CTC and metastatic cancer datasets, we first preprocessed and normalized the datasets using Seurat V3 package [45]. For each dataset, we selected the top 2000 highly variable genes using the FindVariableFeatures function in Seurat V3 with the ‘vst’ selection method. For batch effect elimination, anchors were inferred using the FindIntegrationAnchors function and based on the inferred anchors, the datasets were integrated using IntegrateData function. The integrated data was scaled, and dimension reduction, clustering and visualization steps were followed as described before.

### Cell culture & Reagents

Primary human mesenchymal stem cells (MSCs) were obtained from Lonza (Cat # PT-2501) and MDA-MB-231 human breast cancer cells from NCCS, Pune. MSCs were maintained in complete media consisting of low glucose Dulbecco’s modified eagles medium (DMEM) (Himedia, Cat # AL006A) supplemented with 16% foetal bovine serum (FBS, Thermo Scientific, Cat # 12662029), 1% penicillin-streptomycin (Pen-strep, Sigma, Cat #) and 1% L-glutamine (Thermo Scientific, Cat # 35050061). MDA-MB-231 cells were cultured in DMEM-High Glucose (Himedia, Cat # AL006) with 10% FBS (Himedia, Cat # 1112), 1% pen-strep and 1% L-glutamine. For TGF*β* inhibition experiments, TGF*β* neutralizing antibody (R&D systems, DY240) was added at a concentration of 1.25 μg/ml. Contractility inhibition experiments were carried out by adding the non-muscle myosin II inhibitor blebbistatin (5 μM).

### Fabrication and characterization of PA gels

Polyacrylamide (PA) gels of varying stiffness (0.5, 2, and 5 kPa) were fabricated on 3-APTMS (Sigma) functionalized glass coverslips by combining 40% acrylamide and 2% bis-acrylamide in specific ratios as described previously [81]. Gels were crosslinked by addition of 10% ammonium persulfate (APS) (1:100) and tetramethylethylenediamine (TEMED) (1:1000). After UV crosslinking the gels with Sulfo-SANPAH (Thermo-scientific, Cat # 22589) in 50 mM HEPES buffer (SRL chemicals, Cat # 63732), gels were washed with PBS and treated with 25 μg/ml of Type-I collagen (Thermo-scientific, Cat # A104830) at 4°C overnight. Gels were then washed with PBS to remove unbound collagen and incubated with cell culture media for 30 minutes prior to seeding cells.

### Conditioned media collection

For collection of MDA-MB-231 secreted cancer conditioned media (CCM), cancer cells were seeded on PA gels of 0.5, 2, and 5 kPa stiffness at a seeding density of 12000 cells/cm^2^. Cells were cultured for 48 hrs in complete media. The collected CCM was centrifuged (1500 rpm, 10 minutes) and filtered through a 0.22 μm pore filter prior to use for experiments (Figure 2A). For preparation of CAF-conditioned media (CAFCM), MSC were cultured on different stiffness gels for 7 days in the presence of CCM. After 7 days, differentiated MSCs were washed with PBS and cultured with MSC complete media for next 24 hrs (Fig. 5A). The media was then collected, centrifuged and filtered for further use. The collected conditioned media were stored at −20°C or −80°.

### Design and fabrication of microfluidic devices

For studying CCM-induced MSC chemotaxis, we used our previously designed device [38]. For shear experiments, a straight channel 400 μm in width having 3 ports was designed (Fig. 6G). Shear stress profile in the zone of interest was simulated using COMSOL MultiPhysics (version 5.2) assuming laminar flow conditions. The device designs were printed on to a transparency mask using a printer with resolution 600 dpi and higher. The design was patterned on a silicon wafer by photolithography with SU-8 (2050) photoresist (MicroChem). PDMS devices were then fabricated by pouring PDMS (Sylgard 184) onto the master, and curing them in the oven. After peeling the PDMS substrates, all ports were punched using a 3 mm biopsy punch. Finally, devices were fabricated by bonding the PDMS devices to glass.

### Cell experiments

For MSC chemotaxis experiments, 15 μl media containing 120000 MSCs (prestained with Hoechst 33342) were mixed with different concentrations of 3D collagen (1, 2 and 3 mg/ml) and introduced through one of the channel inlets. Media containing only collagen was introduced through the other inlet. After allowing MSCs to adhere onto the 3D collagen gel (∼6 hrs), stiffness-modulated CCM from was introduced as a chemokine at the opposite channel inlet. For measuring cell motility, time-lapse microscopy was performed for 12 hrs using an inverted microscope equipped with an onstage incubator (Evos FL Auto, Life Technologies). Images were acquired every 15 minutes. Cell motility was quantified using the manual cell tracker plugin in ImageJ (NIH).

For differentiation experiments, MSCs were seeded on PA gels at a density of 800 cells/cm^2^. After 12 hrs of seeding, CCM collected from 0.5, 2, and 5 kPa stiffness were added onto the respective gels in 1:1 ratio (i.e., CCM: fresh media = 1:1) for a duration of 7 days. This combined media was replenished after every 48 hrs. For expression profiling of CAF-associated markers, integrins and MMPs, quantitative real time PCR was performed as described elsewhere [82]. Total RNA was isolated using RNeasy Mini Kit (Qiagen, Cat # 74104). 1 μg of total RNA was used in cDNA synthesis using ProtoScript II Reverse Transcription kit (NEB, Cat # e6560) as per manufacturer’s instructions. Primer details are provided in Supp. Table 1.

For immunostaining, MSCs were fixed with ice-cold 4% paraformaldehyde (pH 7) in the presence of permeabilization buffer (0.5% TritonX-100) for 1 min to remove soluble cytoplasmic proteins. After fixation, cells were washed with cytoskeleton stabilizing buffer (CSB) (60mM PIPES, 27 mM HEPES, 10mM EGTA, 4mM magnesium sulphate heptahydrate, pH 7) three times, and then incubated with blocking buffer (2 %BSA) for 30 min at 4°C. Cells were then incubated with anti-α-SMA antibody (mouse monoclonal, Sigma, Cat# A2547, 1:400 dilution) and anti-LMNA antibody (mouse monoclonal, Sigma, Cat# Ab8980, 1:400 dilution), anti-pMLC (rabbit, CST, cat# 3671S,, 1:400 dilution), anti-Stro-1 (mouse, abcam, cat# ab102969, 1:400 dilution). anti-CD44 (mouse, Novus, cat# 8E2F3, 1:400 dilution), anti-ITGB1 (rabbit, abcam, cat# ab183666, 1:400 dilution) overnight at 4°C diluted in blocking buffer. The following day, cells were washed with CSB thrice and then incubated with respective secondary antibody (Life Technologies, 1:1000 dilution), Phalloidin (Life Technologies, 1:400 dilution) and Hoechst 33342 (Life Technologies, 1:1000 dilution) for 2 hrs at room temperature (RT). Cells were imaged using laser-scanning confocal microscope (LSM 710, Zeiss 40x magnification), Olympus inverted fluorescence microscope(40x, 60x magnification) or Zeiss Spinning disc confocal microscope (20x magnification). Intensity of cells were quantified for multiple cells across 3 independent experiments for each condition using ImageJ (NIH).

For cell proliferation measurements, MSCs cultured on PA gels in the presence and absence of CCM were imaged (15 random images) every alternate day for 7 days. Population doubling was calculated by dividing the average number of cells per frame on a given day by the average number of cells per frame on Day 1. Cell spreading area was determined by manually drawing the outline of individual cells using the polygonal tool in ImageJ (NIH).

For traction force microscopy (TFM) measurements, 5 kPa gels were prepared on 22 × 22 mm^2^coverslips. Once the gels solidified, a thin layer of 25 μl of 5 kPa gel with 1 μm rhodamine fluorescent beads (Fluka, 1:50 dilution) was prepared on previously prepared gel. ECM coating was performed as described in previous section. After 24 hr of cell seeding, the cells were imaged at selected location for phase contrast and Red Flurescence Protein (RFP). Cells were then lysed using Triton-X 100 without moving the plate containing the gels and RFP images were again captured. For calculating the magnitude of traction forces, the Matlab code was used from J. P. Butler [83].

For collagen degradation assays, gels were prepared and coated with DQ collagen overnight at 4°C. After culturing MSCs/CAFs on these substrates for 48 hrs, cells were fixed using 4% PFA, and then stained with Phalloidin and Hoechst 33342, respectively. Fluorescence images (degraded collagen: green, F-actin: red, and nucleus: blue) were captured at 60x magnification. Integrated intensity of degraded collagen per cell was quantified using NIH ImageJ and normalized with respect to that of D1 MSCs.

### Spheroid Experiments

MSCs cultured with CCM on 2 kPa and 5kPa gels were detached using TrypleE, pelleted and resuspended in media. Cancer cells from tissue culture plastic (TCP) dishes were also trypsinized and re-suspended in media. While homospheroids were generated using MDA-MB-231 cells alone (SM), heterospheroids were generated by mixing 2 kPa/5 kPa CAFs and cancer cells in a 1:1 ratio (SMF 2 and SMF 5, respectively) (Fig. 5F). Single spheroids were generated using hanging drop (15 μl) method by incubating 4000 cells in media containing 6.25 μg/ml of rat-tail collagen I (Thermoscientific, Cat # A104830) at 37°C, 5% CO2 for 48 hrs. Cell viability within spheroids was assessed by carrying out live/dead staining using Calcein AM (Life Technologies, Cat # C3099) and propidium iodide (PI, Himedia, Cat # TC-252). For spheroid invasion experiments, spheroids were embedded in 1.5 mg/ml 3D collagen gels by mixing collagen with 10X PBS (phosphate buffered saline) and DMEM at 4°C and kept in incubator for collagen gel formation. After 30 minutes, wells were flooded with media. Images were captured at different time points starting from t = 0 hrs to t = 7 days. The extent of invasion was quantified and normalized with respect to invasion of MDA-MB-231 spheroids (i.e., SM). To assess the spatial position of CAFs in heterospheroids, CAFs were stained with CellTracker red (Thermoscientific, Cat # C34552) and MDA-MB-231 cells with CellTracker green (Thermoscientific, Cat # C2925). Spatial distribution of CAFs/cancer cells was observed 48 hrs after implanting spheroids in collagen gels.

For adhesion experiments, spheroids were seeded on fibronectin (FN) coated glass coverslips. Images were taken at t = 0 to count the number of spheroids seeded and kept for 2 hrs in incubator. Coverslips were washed with PBS after t = 2 hrs to remove non-adherent/weakly-attached spheroids, and then imaged to count the remaining spheroids. Percentage spheroid attachment was calculated across all conditions. For estimating the strength of adhesion, trypsin de-adhesion experiments were performed wherein after 2 hrs of adhesion onto Fn-coated coverslips, spheroids were incubated with Trypsin-EDTA and imaged at 10 sec intervals till the spheroids start to detach, i.e., t = τ* [84, 85]. The time required for detachment, τdetach = (τ ^*^ – 120) mins, corresponds to the time when there is a visible shift observed in the centroid position of the spheroid as seen in the pseudocolor images (pink and blue) in Fig. 6F.

For shear experiments, PDMS devices designed for shear experiments were bonded to glass using plasma oxidation, and were coated with FN overnight at 4°C. Next day, after removing unbound FN with PBS wash, device was loaded with media prior to spheroid seeding. Spheroids were seeded in the middle well and allowed to adhere for 10-12 hrs. Subsequently, media from inlet was flown at a rate of 400 μl/min. τshear corresponds to the time duration for which individual spheroids remained attached under shear. This was determined based on change in the spheroid position before and after flow was started. The position of individual spheroids is shown in pseudocolor (pink and blue) in Fig. 6I.

### Statistical analysis

All statistical analysis was performed using Origin 9.1 with *p* < 0.05 considered to be statistically significant. Based on the normality of data assessed using Kolmogorov-Smirnov normality test, one-way ANOVA/ twoway ANOVA was performed to assess statistical significance, and Fisher post-hoc test was used to compare the means.

## Acknowledgements

The authors thank Centre for Nanoelectronics, IIT Bombay for providing lithography facility and IRCC for Confocal (LSM & Spinning Disc) and AFM Central Facilities. Authors acknowledge financial support from Department of Biotechnology (Govt. of India, Grant # BT/PR7741/MED/32/275/2013) and Department of Science and Technology (Govt. of India, Grant # DST/SJF/LSA-01/2016-17). NS was supported by the Inspire Fellowship from the Department of Science and Technology (Govt. of India, DST/INSPIRE Fellowship/2013/1033).

## Author Contributions

Conceptualization: NS, HZ and SS; Methodology: NS, GB SJ, HZ and SS; Experimental analysis: NS; RNAseq analysis: GB and HZ; Writing: NS, HZ and SS.

