## Supplementary figures and images for "Pre-metastatic niche drives breast cancer invasion by modulating MSC homing and CAF differentiation"

### Supp. Figures

#### Supp Data

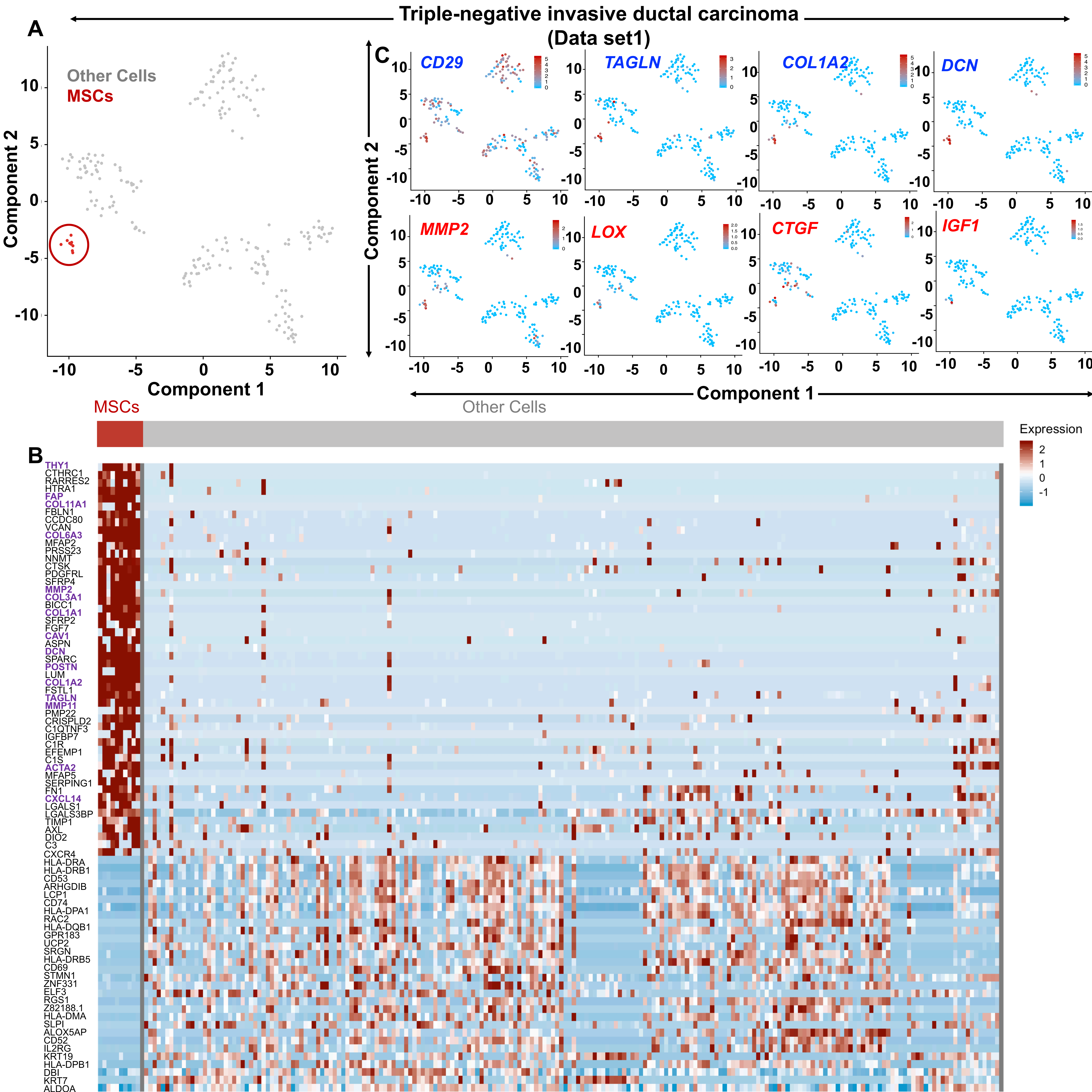

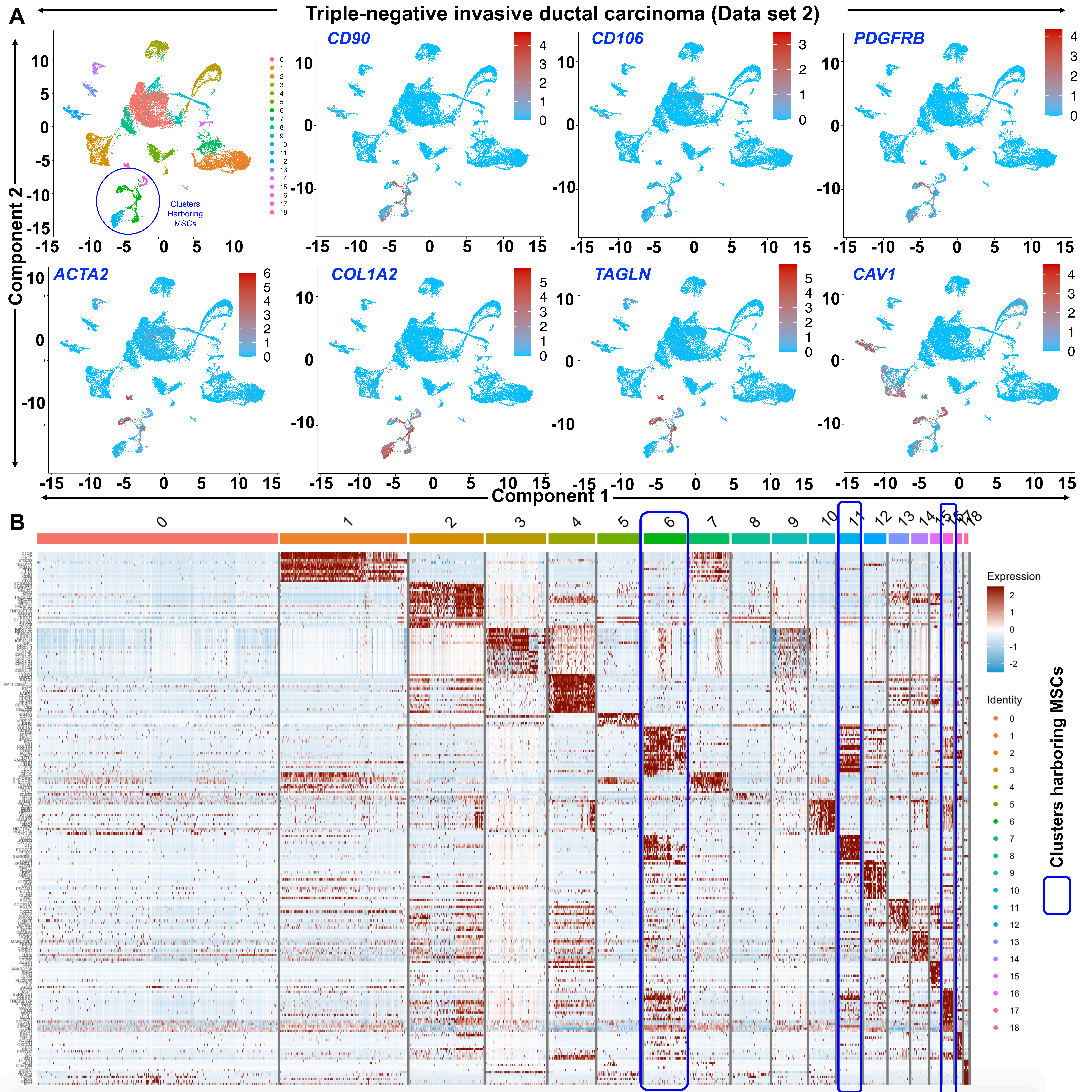

#### Supplementary Fig 3

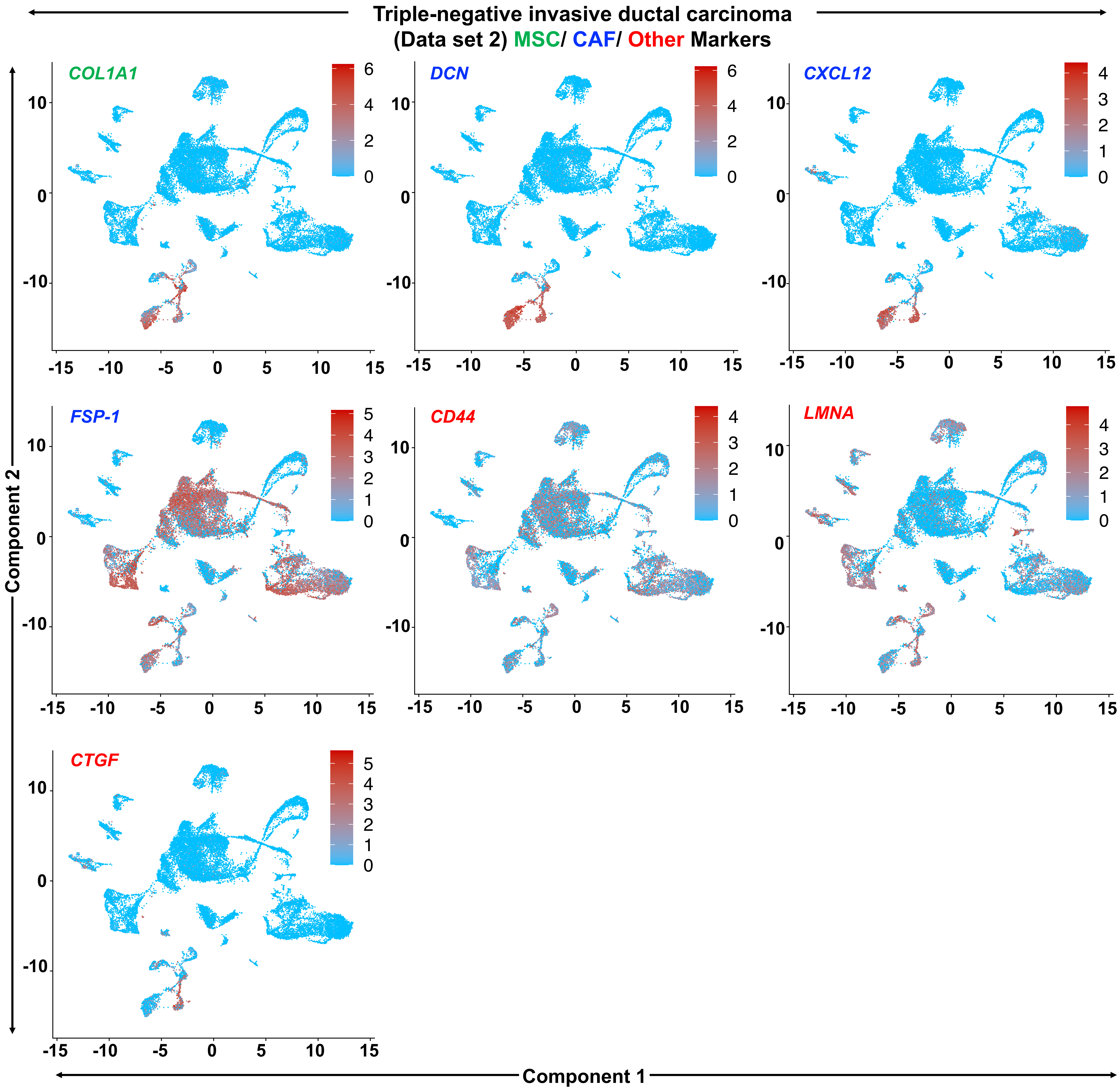

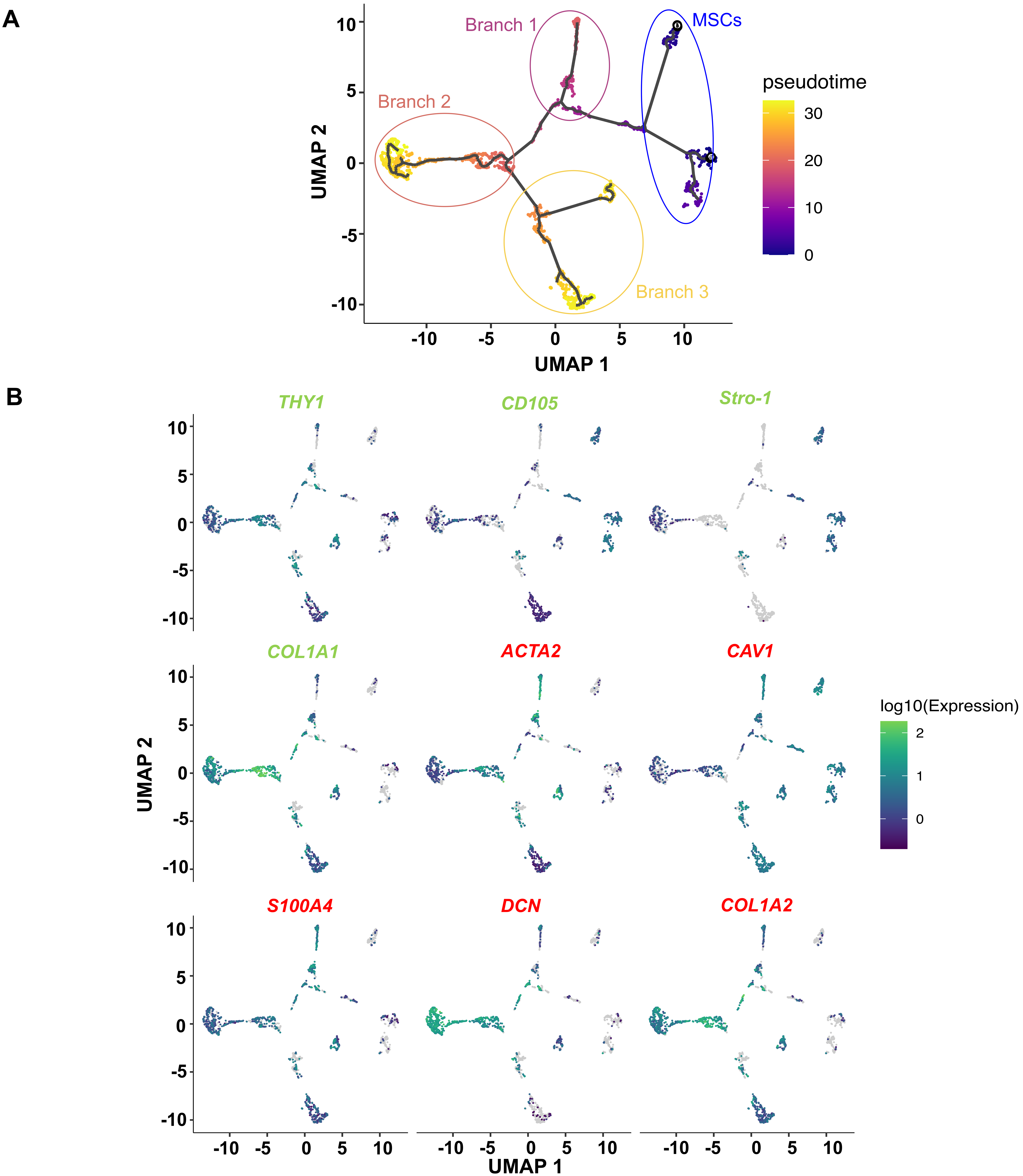

#### Supplementary Fig 5

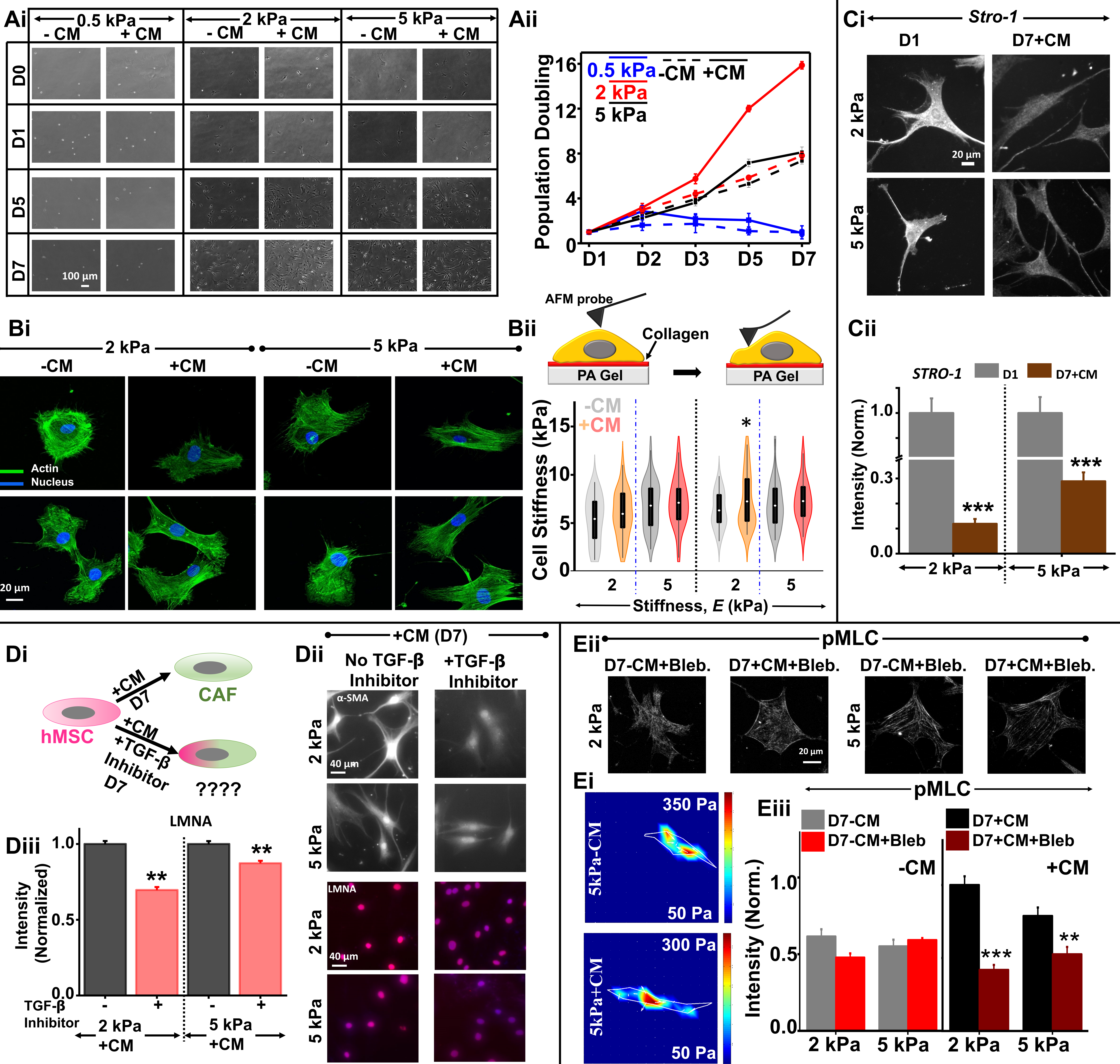

#### Supplementary Fig 6

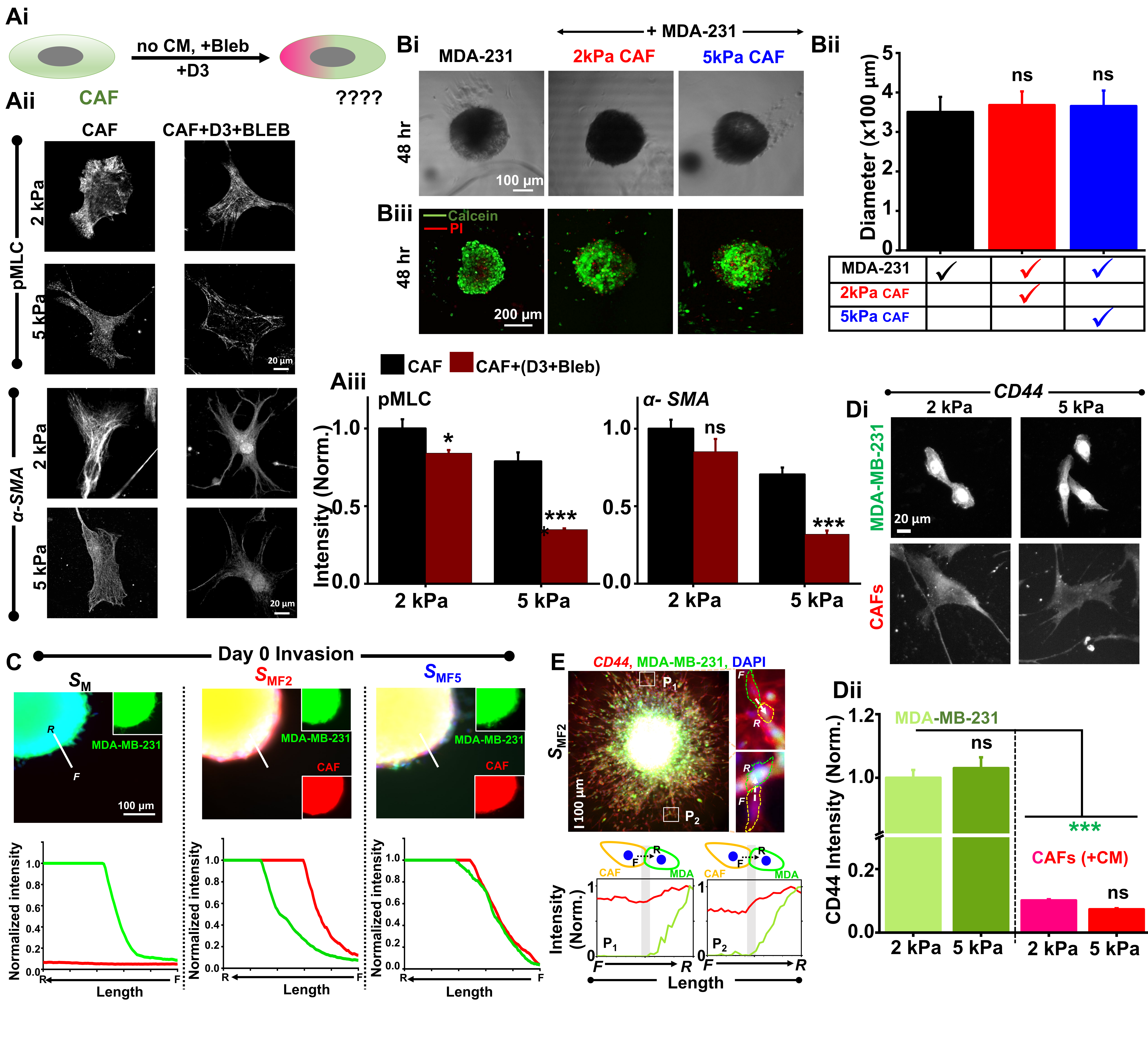

#### Supplementary Fig 7

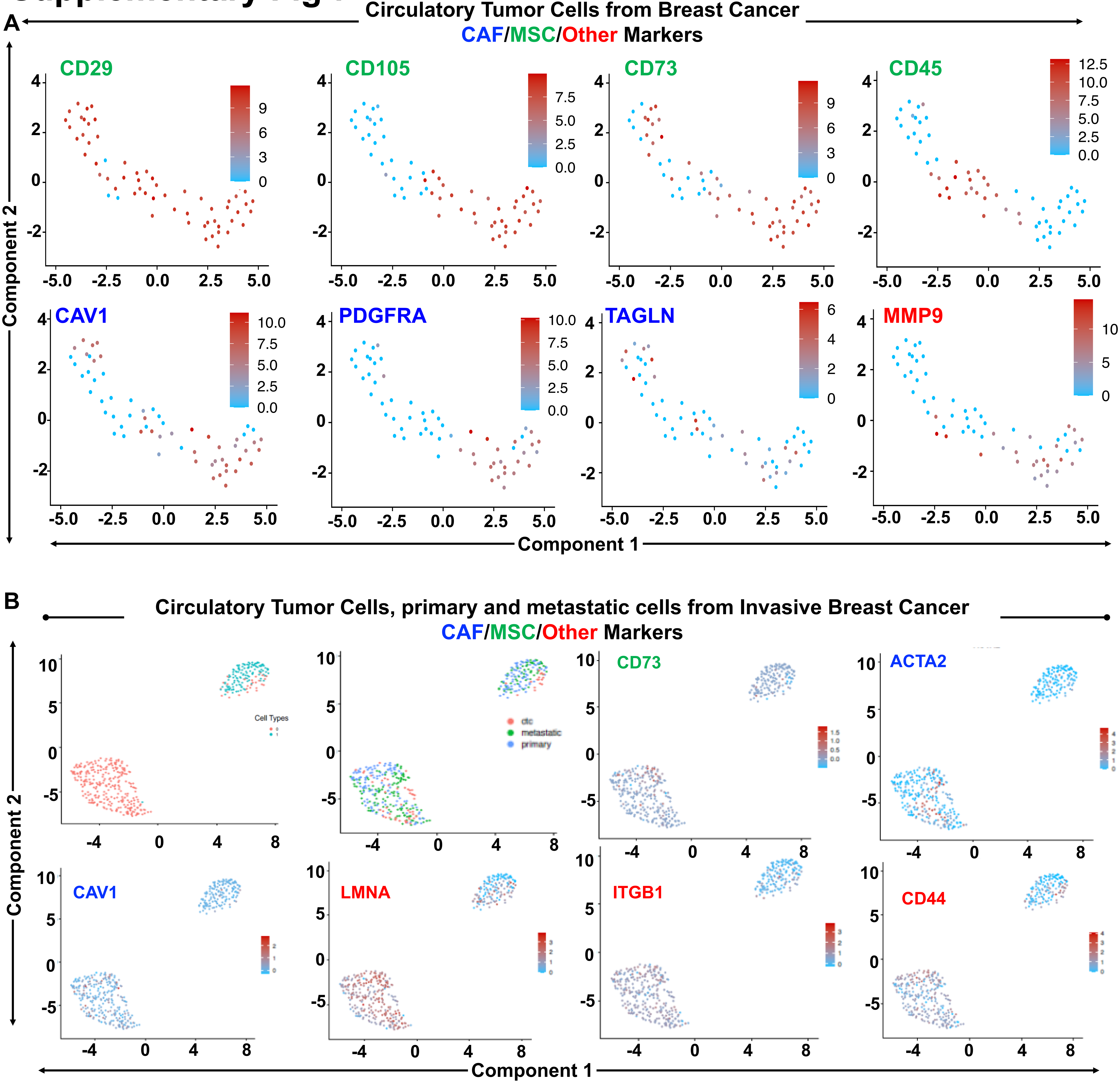
